# The mechanism of cross-talk between histone H2B ubiquitination and H3 methylation by Dot1L

**DOI:** 10.1101/501098

**Authors:** Evan J. Worden, Niklas Hoffmann, Chad Hicks, Cynthia Wolberger

## Abstract

Methylation of histone H3, lysine 79 (H3K79), by Dot1L is a hallmark of actively transcribed genes that depends on monoubiquitination of H2B at lysine 120 (H2B-Ub), and is a well-characterized example of histone modification cross-talk that is conserved from yeast to humans. The mechanism by which H2B-Ub stimulates Dot1L to methylate the relatively inaccessible histone core H3K79 residue is unknown. The 3.0 Å resolution cryo-EM structure of Dot1L bound to ubiquitinated nucleosome reveals that Dot1L contains binding sites for both ubiquitin and the histone H4 tail, which establish two regions of contact that stabilize a catalytically competent state and positions the Dot1L active site over H3K79. We unexpectedly find that contacts mediated by both Dot1L and the H4 tail induce a conformational change in the globular core of histone H3 that reorients K79 from an inaccessible position, thus enabling this side chain to project deep into the active site in a position primed for catalysis. Our study provides a comprehensive mechanism of cross-talk between histone ubiquitination and methylation and reveals an unexpected structural plasticity in histones that makes it possible for histone-modifying enzymes to access residues within the nucleosome core.

## Introduction

Post-translational modifications of histones play a central role in regulating all cellular processes requiring access to DNA. Cross-talk between histone modifications, in which one histone modification regulates deposition of a second (Suganuma and Workman, 2008), provides an additional layer of regulation and specificity. The dependence of histone H3 Lysine79 (H3K79) methylation on monoubiquitination of histone H2B Lysine120 (humans; lysine 123 in yeast) (Briggs et al., 2002; McGinty et al., 2008; Ng et al., 2002c; Steger et al., 2008; Van Leeuwen et al., 2002) is an example of cross-talk that is conserved across eukaryotes. H3K79 mono-, di- and tri-methylation regulates transcription elongation (Kouskouti and Talianidis, 2005; Krogan et al., 2003; Steger et al., 2008), telomeric silencing (Ng et al., 2002a; Takahashi et al., 2011) and the DNA damage response (Giannattasio et al., 2005; Huyen et al., 2004; Wysocki et al., 2005). H3K79 is methylated by human Dot1L (Feng et al., 2002; McGinty et al., 2008) and yeast Dot1 (Van Leeuwen et al., 2002), which are methyltransferase enzymes that attach up to three methyl groups to the lysine *ε* amino group using S-adenosylmethionine (SAM) as the methyl donor (Feng et al., 2002; Van Leeuwen et al., 2002). Human Dot1L is a 1,527 amino acid protein that contains an N-terminal methyltransferase domain that is conserved in yeast Dot1 (Sawada et al., 2004), and a C-terminal extension that interacts with proteins that direct Dot1L to specific genomic loci (Kuntimaddi et al., 2014). Mislocalization of Dot1L due to gene fusions between Dot1L-interacting proteins and the histone H4 methyltransferase, Mixed Lineage Leukemia (MLL) can lead to hypermethylation of H3K79 and constitutive gene expression at ectopic loci, which is a causative factor for around 40% of Mixed Lineage Leukemias (MLL) (Bernt et al., 2011; Chen and Armstrong, 2015; Guenther et al., 2008; Krivtsov et al., 2008). Despite multiple studies, the mechanism by which H2B-Ub stimulates Dot1L to methylate H3K79 remains unknown.

The dependence of H3K79 methylation on prior monoubiquitination of histone H2B (H2B-Ub) has been established both *in vivo* and *in vitro* (Briggs et al., 2002; McGinty et al., 2008; Ng et al., 2002b). Methylation of H3K79 in yeast is abolished in strains lacking either the E2 or E3 that ubiquitinate H2B-K123, Rad6 and Bre1, or in strains in which histone H2B bears an arginine in place of K123 (Briggs et al., 2002; Ng et al., 2002b; Wood et al., 2003). Knockdown of human RNF20 or RNF40, which form the RNF20/40 heterodimeric E3 ligase that ubiquitinates H2BK120, leads to a decrease in H3K79 methylation in human cells (Wang et al., 2013; Zhu et al., 2005). In vitro, Dot1L is directly stimulated by the presence of H2B-Ub in mononucleosomes (McGinty et al., 2008), which increases the apparent catalytic rate of H3K79 methylation (McGinty et al., 2009). Both Dot1L and yeast Dot1 require a nucleosomal substrate and are not active on isolated histone H3, suggesting that these methyltransferases may recognize other portions of the nucleosome (Feng et al., 2002; Van Leeuwen et al., 2002). In another layer of regulation, the H4 tail has been implicated in contacting Dot1/Dot1L and is required for H3K79 methylation by both human Dot1L (McGinty et al., 2009) and yeast Dot1 (Altaf et al., 2007; Fingerman et al., 2007). Deletions and point substitutions in H4 residues R17-H18-R19 dramatically reduce methylation of H3K79 in yeast (Altaf et al., 2007; Fingerman et al., 2007) and abolish in vitro activity of both yeast Dot1 (Fingerman et al., 2007) and human Dot1L (McGinty et al., 2009). Although structures of the Dot1 (Sawada et al., 2004) and Dot1L (Min et al., 2003; Richon et al., 2011) catalytic domains have been determined, it is still not known how the enzyme recognizes ubiquitin or contacts other portions of its nucleosomal substrate. Due to the limited accessibility of the H3K79 side chain, which is oriented along the nucleosome surface (Lu et al., 2008), it has not been possible to construct a plausible model of Dot1L bound to its nucleosomes in which the K79 side chain can enter the active site and attack the bound SAM cofactor (Min et al., 2003).

We report here cryo-EM structures of Dot1L bound to ubiquitinated nucleosomes that reveal the molecular basis for cross-talk between histone H2B ubiquitination and H3K79 methylation and uncover unexpected allostery in both enzyme and substrate that stimulates catalysis. We have captured a complex of Dot1L bound to H2B-Ub nucleosome and SAM in a catalytically competent state, which was determined at a resolution of 2.96 Å. The structure of Dot1L bound to H2B-Ub nucleosomes in a poised state, in which the active site is too far from H3K79 for catalysis, was determined at a resolution of 3.95 Å. The structures reveal that Dot1L contains structural elements that contact the H2B-conjugated ubiquitin and the nucleosome H2A/H2B acidic patch, which establish a pivot point about which Dot1L can rotate. In the active complex, the histone H4 tail binds to a previously uncharacterized groove in Dot1L, tethering Dot1L to the opposing edge of the nucleosome and precisely positioning the catalytic center above H3K79. Remarkably, a conformational change within the globular core of histone H3 repositions K79 from an inaccessible conformation to one that inserts the side chain into Dot1L’s active site in a position to initiate the methyl transfer reaction. The altered conformation of histone H3 is stabilized by contacts with both the H4 tail and with Dot1L, which itself undergoes conformational changes that accommodate binding of the H4 tail and enclose the substrate lysine in a hydrophobic channel. The structure accounts for all prior observations regarding the role of ubiquitinated nucleosomes in activating Dot1L and the yeast homologue, Dot1, and suggests new avenues for designing inhibitors that are specific for the Dot1L catalytic domain.

## Results

### Structural overview of Dot1L bound to nucleosomes containing H2B-Ub

We determined cryo-EM structures of Dot1L bound to nucleosomes in which ubiquitin was conjugated to histone H2B-K120 via a non-hydrolyzable dichloroacetone linkage (Morgan et al., 2016). For the poised state structure, we formed complexes between Dot1L and H2B-Ub nucleosomes in the presence of an analogue of the SAM methyl donor (see Methods), although we see no bound cofactor in our maps. To trap Dot1L in the active state, we formed complexes between Dot1L, SAM, and H2B-Ub nucleosomes containing histone H3 in which K79 was replaced with norleucine (Nle) (see Methods), a non-native amino acid that has been shown to increase the affinity of methyltransferases for peptides in a SAM-dependent manner (Brown et al., 2014; Jayaram et al., 2016; Van Hest et al., 2000). Norleucine has the same number of aliphatic carbons as lysine but lacks the charged *ε* amino group. The structure of the complex in the poised state was determined to a resolution of 3.90 Å and structure of a complex in the active state, in which H3K79 extends into the Dot1L active site, was determined to a resolution of 2.96 Å (Figures 1a,b, S1-3 and Table 1). Two classes of active state complexes were identified: one (36% of particles) in which Dot1L is bound to both faces of the nucleosome (Figures 1a S2) and one (34% of particles) in which one Dot1L is bound to the nucleosome (Figure S2). All complexes contain a fragment of human Dot1L (residues 2-416) that contains the catalytic domain and a C-terminal DNA binding region, and that can be fully stimulated by nucleosomes containing H2B-Ub (McGinty et al., 2008; Min et al., 2003). In both structures, only Dot1L residues ~4-332 are visible in the maps, indicating that the C-terminal DNA binding region of Dot1L is either disordered or highly mobile.

**Figure 1:**
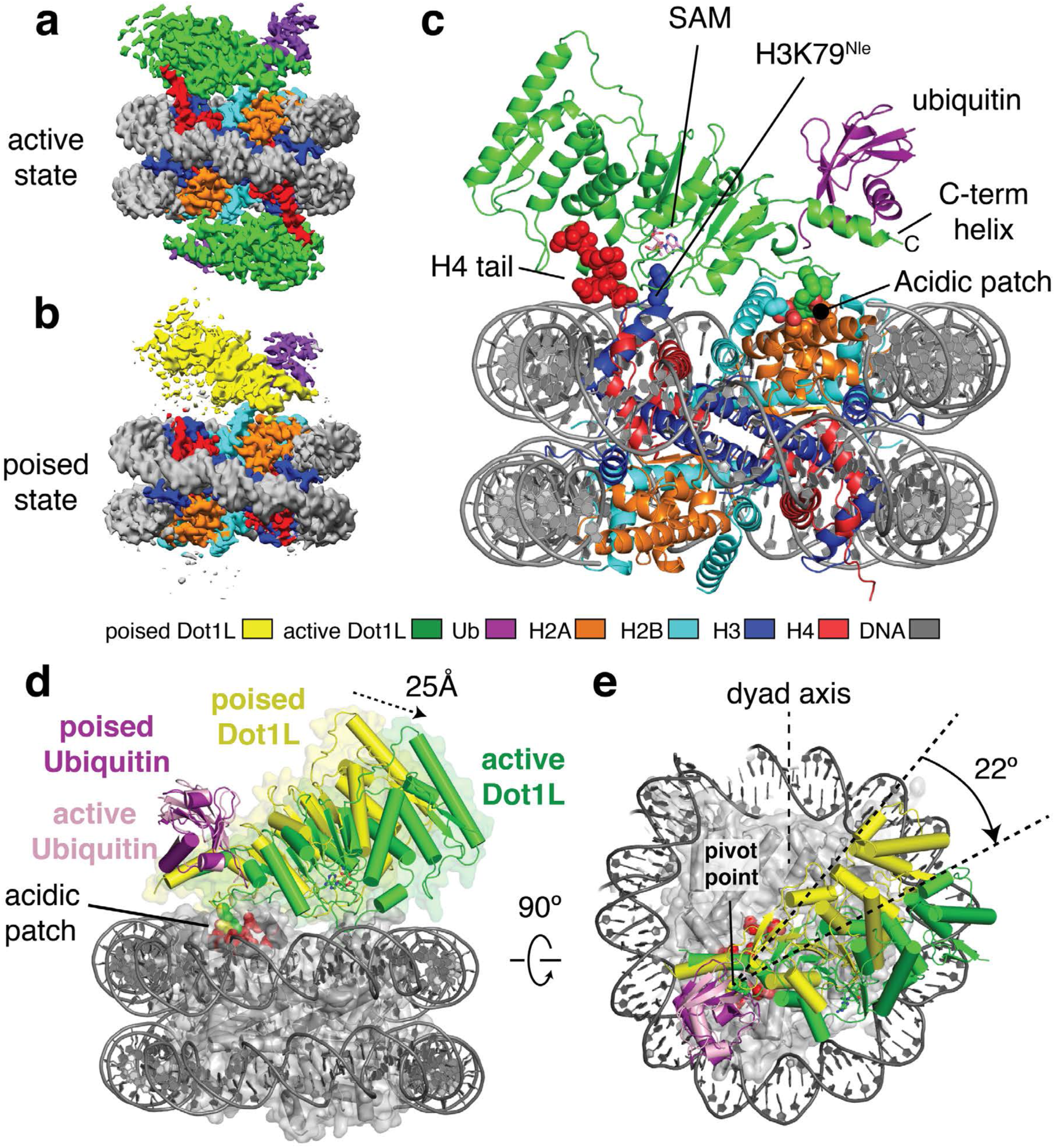
Structure of the Dot1L complex with the H2B-Ubiquitin nucleosome. **a**, Cryo-EM reconstruction of the active state complex. **b**, Cryo-EM reconstruction of the poised state complex. **c**, Atomic model of the active state complex between Dot1L and the H2B-Ub nucleosome looking down the dyad axis. The H4 tail, H3K79^Nle^ and the acidic patch are depicted as spheres and the SAM cofactor is depicted in stick representation. **d**,**e** Dot1L movements from the poised state to the active state shown as from the side and top, respectively. Poised state and active state Dot1L are depicted as yellow and green cylinders and the histone octamer is shown as a transparent gray surface. Acidic patch residues are shown as red spheres. Black arrows indicate movements made by Dot1L in the switch to the active state.

In both the active and poised state complexes, the Dot1L C-terminus contacts ubiquitin and the nucleosome acidic patch formed by histones H2A and H2B, anchoring Dot1L to one edge of the nucleosome (Figures 1c,d and S6). As compared to the poised state, Dot1L in the active state is rotated clockwise by 22° about the ubiquitin and pivots down towards the nucleosome face by 25 Å at its N-terminus (Figure 1d,e). The active state is further stabilized by binding of histone H4 tail residues 15-19 in a previously unknown groove in the N-terminal portion of Dot1L (Figure 1c). The greater number of contacts between Dot1L and the nucleosome in the active complex appear to stabilize the enzyme, as evidenced by higher map resolution and well-resolved density throughout the catalytic domain (Figure S4). Importantly, the H3K79^Nle^ side chain projects into the active site immediately adjacent to the SAM methyl group (Figure 1c). This close approach is facilitated by an unprecedented conformational change in the histone fold of H3 that shifts the K79 side chain from an inaccessible orientation, enabling it to enter the Dot1L active site in a position poised for catalysis. With the exception of conformational changes in two Dot1L loops described below, the structure of Dot1L is essentially identical to that previously reported (Figure S5a,c) (Min et al., 2003; Richon et al., 2011).

The poised structure represents a catalytically incompetent state in which Dot1L cannot methylate H3K79 (Figures 1b, and S6) and likely exists directly before or after methyl-transfer. Dot1L contacts ubiquitin and the acidic patch in essentially the same manner as seen in the active state structure (Figures S5d and S6a-d). In the poised state, the N-terminal portion of the Dot1L catalytic domain does not contact the nucleosome and the Dot1L active site is ~21 Å away from H3K79 (Figure S6e). The relatively weak density for the N-terminal portion of Dot1L, as compared to the C-terminal portion that contacts ubiquitin and the nucleosome (Figures 1b and S6a), suggests that Dot1L can occupy a continuum of conformations when the N-terminal part of the enzyme does not contact the nucleosome. Due to the higher resolution and better density of the active state structure, all images and discussion below focuses on the active state structure unless otherwise noted.

### Contacts with ubiquitin and the H2A/H2B acidic patch orient Dot1L on the nucleosome

The structure reveals that Dot1L contains a ubiquitin-binding hydrophobic cradle comprising the C-terminal helix (residues 320-330) of the catalytic domain and a loop between β-strands 8-9, which forms a convex surface that binds ubiquitin (Figures 2a,b and S5d). The interface between Dot1L and ubiquitin buries 637Å^2^ of surface area. While nearly all ubiquitin-binding motifs bind to a canonical patch centered on L8 and I44 of ubiquitin (Komander and Rape, 2012), Dot1L contacts a hydrophobic patch comprising ubiquitin residues I36, L71, and L73 (Figure 2b). Residues L322 and F326 in the ubiquitin-binding helix abut ubiquitin residues I36 and L71, while Dot1L residues L284, I290 and L322 cradle ubiquitin residue L73, which is located at the start of the ubiquitin C-terminal tail (Fig. 2b).

**Figure 2:**
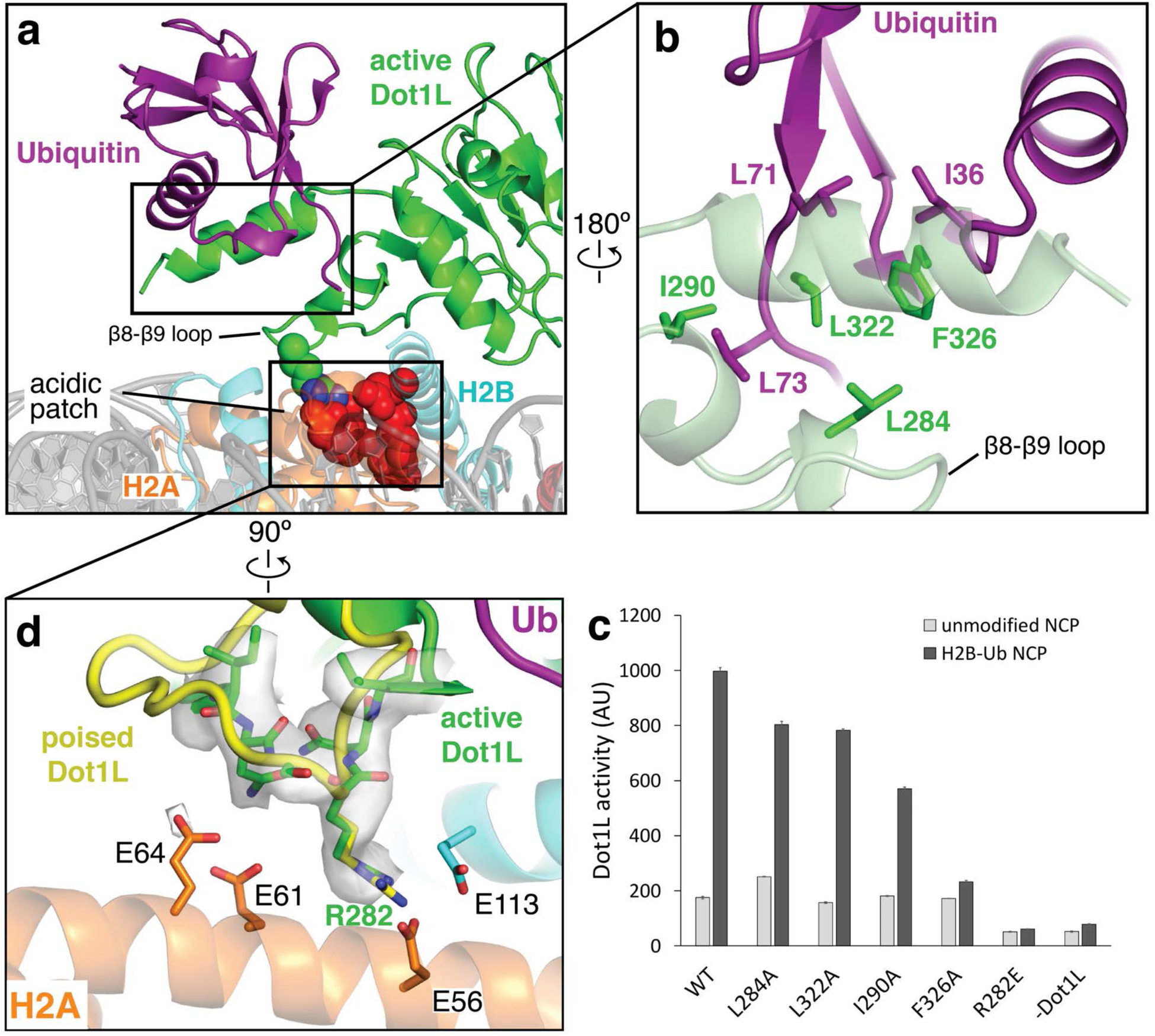
Dot1L interactions with Ubiquitin and the acidic patch. **a**, Overview of the interaction between Dot1L, Ubiquitin and the nucleosome in the active state structure. The nucleosome is depicted as a semi-transparent cartoon. Residues important for Dot1L interaction with the nucleosome acidic patch are indicated and depicted as spheres. **b**, Detailed view of the contacts made between Dot1L and ubiquitin. Dot1L is depicted as a semi-transparent green cartoon. Important residues at the Dot1L-Ubiquitin interface are shown as sticks. **c**, Endpoint H3K79 methylation activity assays using Dot1L mutants with either unmodified or H2B-Ub nucleosomes. Error bars correspond to the standard deviation of 3 replicate experiments. **d**, Detailed view of interactions between Dot1L and the H2B/H2A acidic patch. Residues at the interface are depicted as sticks and the EM density for Dot1L in the active state is shown as a semi-transparent gray surface. A superimposed poised state Dot1L is depicted in yellow.

The observed interactions with ubiquitin explain previous findings that a hotspot involving ubiquitin residues L71 and L73 is critical for ubiquitin-dependent activation of Dot1L (Holt et al., 2015; Zhou et al., 2016), rather than the canonical L8/I44 patch contacted by the majority of ubiquitin-binding proteins (McGinty et al., 2009) (prior findings summarized in Table 2). To probe the contributions of individual Dot1L residues to ubiquitin-dependent stimulation, we generated alanine substitutions in Dot1L residues that contact ubiquitin and assessed the ability of the mutant proteins to be stimulated by H2B-Ub in vitro (Figure 2c). Whereas alanine substitutions L284A, I290A, L322A and F326A did not impair methylation of unmodified nucleosomes, all of these Dot1L substitutions impaired simulation by H2B-Ub-containing nucleosomes (Figure 2c). The most severe defects resulted from an alanine substitution at F326, which was not stimulated at all by nucleosomes containing H2B-Ub. F326 is a highly conserved Dot1L residue (Figure S7) that contacts I36 of ubiquitin, and which is clearly resolved in our maps (Figures 2b and S4c). An alanine substitution at Dot1L residue I290, which is in van der Waals contact with ubiquitin residue L73, reduced stimulation by H2B-Ub to an intermediate level, while the L284A and L322A substitutions had modest effects. Taken together, the effects of both Dot1L (Figure 2b,c) and ubiquitin (Holt et al., 2015) mutations on the ability of H2B-ubiquitinated nucleosomes to stimulate Dot1L confirm the presence of a new ubiquitin binding element that is specific for the hydrophobic base and tail region of ubiquitin.

The pivot point for the switch between the poised and active state is centered on Dot1L residue R282, which contacts the nucleosome acidic patch in both states (Figures 2d and S6d). R282 projects into the H2A/H2B acidic patch and interacts with residues H2A-E56 and H2B-E113 (Figures 2d, and S6c.d). Density for the Dot1L R282 side chain is well-resolved in both active (Figure 2d) and poised states (Figure. S6d), reflecting the stability of this contact. The interaction mediated by R282 is reminiscent of other nucleosome binding proteins that employ an “arginine anchor” to interact with the nucleosome acidic patch (Figure S8) (Armache et al., 2011; Barbera et al., 2006; Makde et al., 2010; McGinty et al., 2014). However, whereas all reported arginine anchor-containing proteins bind to the acidic patch near H2A D90-E92, the Dot1L R282 interaction occurs at a distinct location in the acidic patch (Figure S8). Consistent with its buried position in the poised and active states, a Dot1L R282E substitution completely abrogated the ability of Dot1L to methylate both unmodified and ubiquitinated nucleosomes (Figure 2c).

### The H4 tail binds Dot1L and orients the active site over H3K79

In the active complex, we unexpectedly found that a stretch of basic residues in the tail of H4, 15-19, binds within a groove in the N-terminal portion of Dot1L, tethering the enzyme to the opposite edge of the nucleosome (Figure 3a,b). These H4 residues are not ordered in the poised structure (Figure S9a) and are either unstructured or closely associated with the DNA in most nucleosome structures (Figure S9b). The Dot1L loop 22-32 moves by ~5 Å as compared to the Dot1L crystal structure (Min et al., 2003), opening a cleft in Dot1L that enables the H4 tail to bind (Figures 3b and S9c-d), burying 770Å^2^ of surface area. The sidechain of histone H4 R19 is well ordered in the structure (Figure 3b) and is in a position to interact with the backbone atoms of H3 residues T80, Q76, and K79^Nle^ (Figure 3c). The sidechain of H18 is positioned close to Dot1L residues N126 and S304 and may form hydrogen bonds with these residues (Figure 3c). Histone H4 residue R17 is oriented in a position to project into a deep acidic pocket containing highly conserved residues E138 and Y115 (Figures 3b,c,d and S7). The position of R17 is model built based on clear density for the b carbon and spatial constraints in the acidic pocket. Concurrent binding of the H4 tail and ubiquitin to opposing ends of Dot1L position the enzyme with its catalytic site directly over H3K79, thus ensuring specificity for this residue.

**Figure 3:**
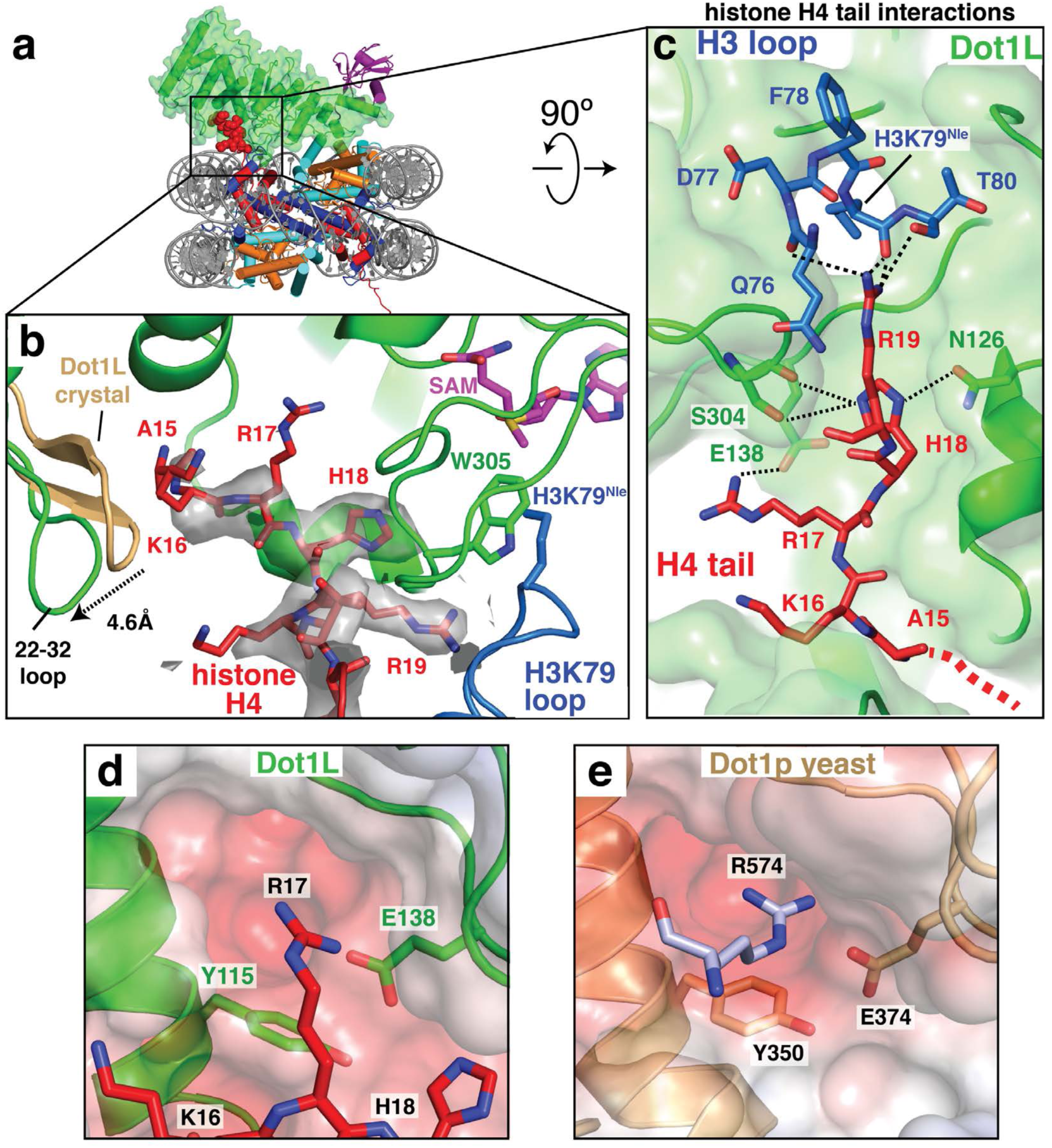
H4 tail interactions with Dot1L. **a**, Overview of the active Dot1L structure. Dot1L shown as a transparent green surface, ubiquitin as purple ribbon, H4 tail in red spheres. **b**, H4 tail (red) interaction with the Dot1L binding groove. EM density for the H4 tail is shown as a semi-transparent gray surface. The 22-32 loop from the crystal structure of Dot1L alone (PDB ID 1NW3) is colored tan. **c**, H4 tail (red) interactions between Dot1L (green) and the H3K79 loop (blue). The surface of Dot1L is shown in semi-transparent green. Potential hydrogen bonding or van der Waals interactions are shown as black dashed lines. Red dashed lines illustrate the direction of the H4 mainchain in the Dot1L binding groove. **d**, Modeled position of H4 R17 binding in the Dot1L acidic pocket. Conserved residues in the binding pocket are depicted as sticks and the surface of Dot1L is shown and colored according the electrostatic potential. **e**, Close up view yeast Dot1p (PDB ID 1U2Z) showing arginine from a neighboring Dot1p molecule in the crystal bound in the conserved acidic pocket. Electrostatic surface potential shown as in **d**. Electrostatic potential in **d, e**, was calculated using the APBS tool.

The observed interactions between Dot1L and the H4 tail explain previous studies showing that H4 residues R17-H18-R19 are required for H3K79 methylation by both Dot1L (McGinty et al., 2009) and yeast Dot1 (Altaf et al., 2007; Fingerman et al., 2007). *In vitro* experiments have shown that a Dot1L R17A/R19A mutant is defective in methylating both ubiquitinated and unmodified nucleosomes (McGinty et al., 2009). Furthermore, deletion of the RHR motif or alanine substitutions at H4 residues R17 or R19 completely eliminate H3K79 di- and tri-methylation by yeast Dot1 and severely reduce mono-methylation of H3K79 both *in vivo* and *in vitro* (Altaf et al., 2007; Fingerman et al., 2007). Due to conservation in the Dot1L catalytic domain and histone residues between yeast and human, these H4 residues are expected to form similar interactions in yeast Dot1p that we observe in the structure of Dot1L. Remarkably, the structure of yeast Dot1 (Sawada et al., 2004) contains a fortuitous interaction in the crystal lattice that appears to mimic binding of H4 R17 to Dot1L (Figure 3e). In the yeast Dot1 crystal structure, an arginine (R574) from the flexible C-terminal tail of a neighboring Dot1 protein in the crystal lattice docks in the conserved acidic pocket of yeast Dot1p (Figure 3e). This interaction had been proposed to reflect a potential interaction with histone H3 (Sawada et al., 2004), but instead likely reflects binding of the H4 tail as observed in our structure of Dot1L. Consistent with the importance of the acidic tunnel in which this arginine binds, substitutions in yeast Dot1 residues Y350 and E374 (Figure 3d), which correspond to human Dot1L Y115 and E138, dramatically decrease Dot1 methyltransferase activity (Sawada et al., 2004). The structure of the active Dot1L complex, combined with the effects of histone H4 mutations on both Dot1L and yeast Dot1, explains the importance of H4 tail interactions in Dot1L activity.

### An induced conformational change in histone H3 orients H3K79 for catalysis

Previous attempts to model a catalytically competent complex by docking the Dot1L crystal structure on the nucleosome were unsuccessful because no orientation of Dot1L can accommodate the H3K79 substrate lysine without significant steric clashes with the core histones (Min et al., 2003). These clashes occur because, in the poised state structure and in other nucleosome structures, H3K79 is in an inaccessible conformation with its side chain projecting parallel to the nucleosome surface (Figures 4a,b and S10a) and is unable to adopt a rotamer capable of entering the Dot1L active site. However, binding of Dot1L in the active state induces a dramatic conformational change in the H3K79^Nle^ loop that reorients both the backbone and side chain of H3K79^Nle^ by ~90°, enabling the side chain to point away from the nucleosome surface and into the Dot1L active site (Figures 4a-e and 5a). The result of this transition is that the H3K79 lysine sidechain moves up and away from the nucleosome surface, which moves the *ε* amino group by ~10 Å (Figure 4b). A modeled H3K79 sidechain based on the position of the aliphatic side chain in the H3K79^Nle^ substitution (Figure 4e) shows that, in the active state, the *ε*-amino group of lysine comes within 3.0 Å of the SAM methyl donor and is positioned for nucleophilic attack on the SAM methyl group (Figure 5a). Importantly, the conformational change in the loop containing H3K79^Nle^ is not due to substitution of lysine with norleucine, since the class of active state complexes containing nucleosomes with just a single Dot1L bound (Figure S2) show that H3K79^Nle^ on the unoccupied face of the nucleosome is in the normal, inaccessible conformation (Figure S10c-e). The unprecedented change observed here in the core histone fold has, to our knowledge, never been observed before in intact nucleosomes and has important implications for how other histone-modifying enzymes may access inaccessible or buried histone residues.

**Figure 4:**
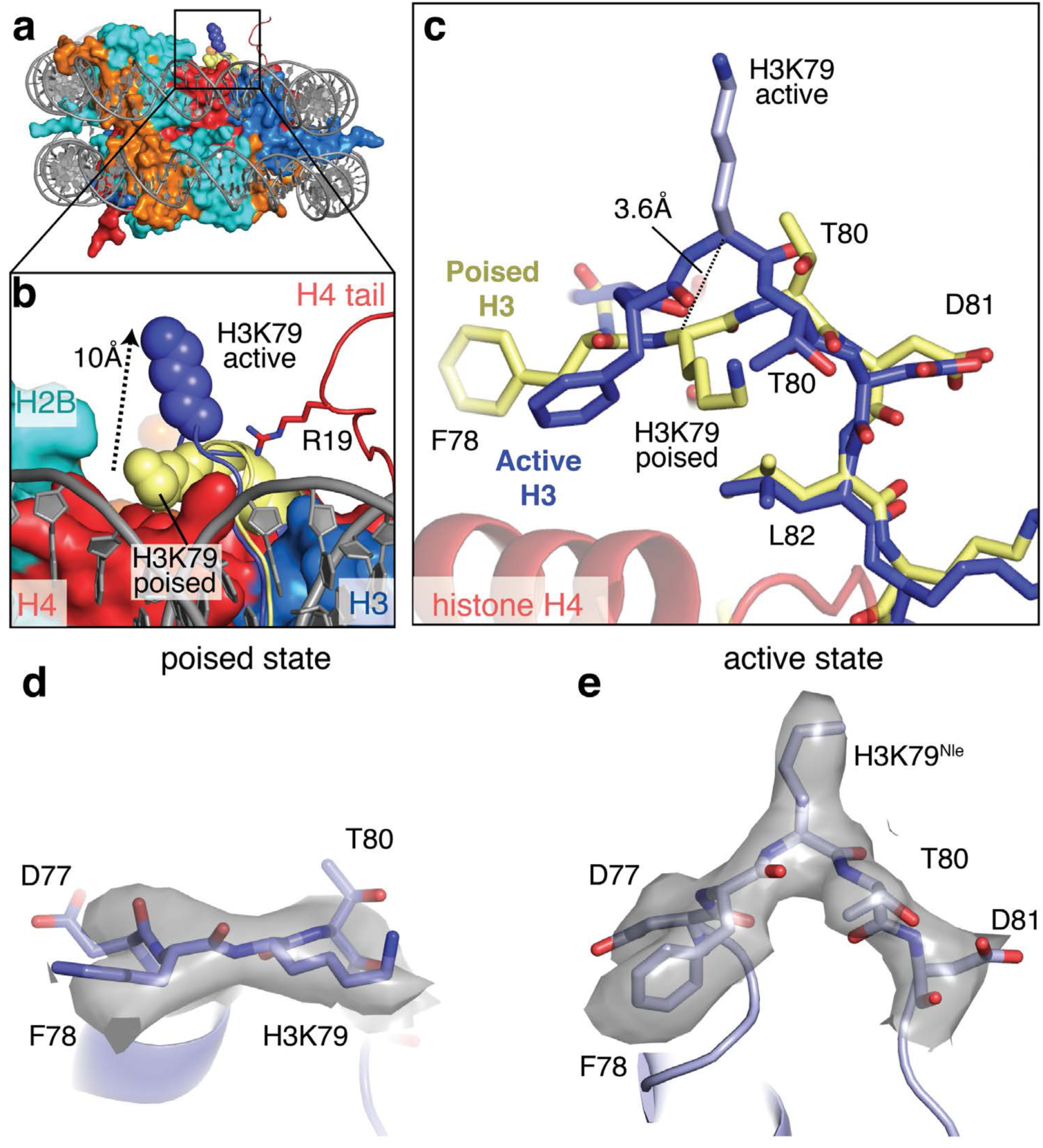
Conformational change in histone H3. **a**, Superimposition of the active state and poised state nucleosomes. The active state histone octamer is depicted in surface representation and colored as in Figure 1. The H3K79 loops from the active state (blue) and poised state (yellow) structures are shown as cartoons and H3K79 is shown as spheres. **b**, Close up view of the active state and poised state H3K79 loops from a. The H3K79 sidechain moves ~10 Å (measured from *ε*-amino groups) from the poised state to the active state. **c**, Superimposition of the histone H3K79 loop from the active and poised state structures showing the conformational change that occurs in the transition from the poised state to the active state. The H3K79 loop in both states is depicted in stick representation. The poised state H3 loop is colored yellow and the active state H3 loop is colored blue. In the transition from the poised state to the active state the H3K79 backbone moves up by 3.6 Å (measured from the Cα carbons of H3K79) and rotates by an angle of ~90°. **d**, The H3K79 loop in the poised state is shown in stick representation and the experimental EM density is depicted as a semi-transparent gray surface. **e**, The H3K79 loop in the active state is shown in stick representation and the experimental EM density is depicted as a semi-transparent gray surface.

The tail of histone H4 and the Dot1L loops containing F131 and W305 play a critical role in stabilizing the active conformation of histone H3 (Figure 5a,b). The loop containing the H3K79 residue is sandwiched between Dot1L residues on one side and H4 tail residues on the other, thus helping to stabilize the altered conformation of histone H3 (Figure 5a,b). Histone H4 R19 hydrogen bonds with backbone atoms of H3 residues K79, T80 and Q76 on one side, while Dot1L residue F131 is in van der Waals contact with the side chain of H3 residue, T80, on the other side of the altered H3 segment (Figures 3c and 5a,b). Upon transition from the poised state to the active state, the highly conserved F131 residue inserts into the same location on the nucleosome that is occupied by the H3K79 side chain in the poised state, further favoring the H3 conformation seen in the active state (Figures 5b and S7). Additional van der Waals contacts between W305 and the aliphatic side chain of H3K79^Nle^ further stabilize the lysine and orient it towards the SAM methyl group. Both F131 and W305 are invariant and reside in loops that are disordered in the poised state but become structured in the transition to the active complex (Figures 5a and S7). Previous studies have determined that the flexibility of these loops can be modulated by binding to SAM and SAM-like inhibitors (Yu et al., 2012). However, we now find that the W305 and F131 loops only become fully structured in the context of the active, nucleosome-bound Dot1L complex. To test the importance of these residues for activity, we assessed the methyltransferase activity of Dot1L mutants bearing an F131A or W305A substitution. As shown in Figure 5d, these substitutions completely abrogated Dot1L activity on both H2B-ubiquitinated and unmodified nucleosomes, consistent with an important role for these contacts in histone methylation.

**Figure 5:**
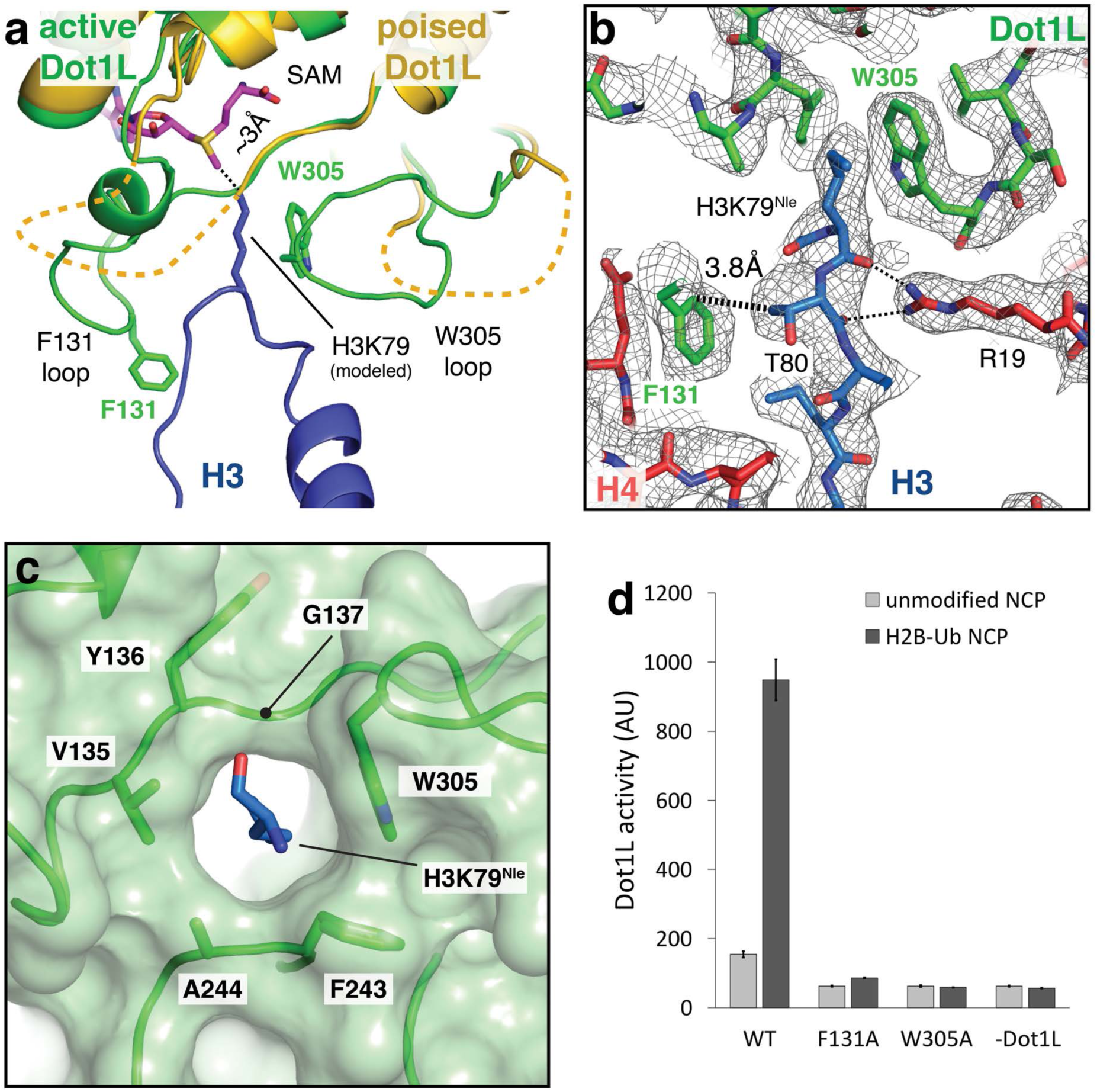
Formation of the Dot1L active site enclosure. **a**, Superimposition of active state (green) and poised state (yellow) Dot1L. The H3K79 loop from the active state structure is shown as a blue cartoon and the modeled H3K79 sidechain is shown in stick representation. The disordered F131 and W305 loops from the poised state structure are depicted as yellow dashed lines. The *ε*- amino group of lysine comes within 3Å of the SAM methyl donor. **b**, Close up view of the Dot1L active site enclosure with H3 (blue), Dot1L (green) and H4 (red) shown in stick representation. Important residues in the Dot1L active site are labeled and the sharpened, experimental EM density is shown as a gray mesh. A van der Waals contact between Dot1L F131 and H3 T80 is shown as thick a dashed black line and the hydrogen bonds between R19 and the H3K79 loop are shown as thin black dashed lines. **c**, Formation of the Dot1L H3K79 lysine binding channel. Dot1L is depicted as a green cartoon surrounded by a semi-transparent green surface. H3K79^Nle^ is shown as blue sticks. **d**, Endpoint H3K79 methylation activity assays using Dot1L mutants with either unmodified or H2B-Ub nucleosomes. Error bars correspond to the standard deviation of 3 replicates.

The repositioning of W305 from a disordered loop completes formation of a narrow, hydrophobic lysine-binding tunnel in which H3K79 inserts (Figure 5c). The other hydrophobic residues that surround the aliphatic portion of the side chain are V135 and F243, thus completely surrounding H3K79 in hydrophobic environment. All three of these hydrophobic residues are invariant (Figure S7). Interestingly, the residue corresponding to W305 in yeast Dot1 is in the active conformation even in the absence of a bound substrate lysine (Figure S10b), thus preforming the tunnel in the yeast enzyme. The methyl group of the bound SAM cofactor caps the end of the tunnel, further encapsulating the attacking H3K79 side chain in a completely hydrophobic environment.

The very hydrophobic lysine-binding tunnel, capped by the hydrophobic methyl group of the bound SAM cofactor, suggests a resolution to the long-standing problem of how Dot1L activates H3K79 for attack on the SAM methyl group, which is the first step in the methylation reaction. The lysine must first be deprotonated; however, previous studies have failed to identify an active site residue that deprotonates the lysine (Min et al., 2003). We see no candidate ionizable residues in the active Dot1L complex. Instead, we speculate that the hydrophobic nature of the tunnel and narrow dimensions that exclude water may be sufficient to lower the pK_a_ of the lysine sufficiently to remove the proton from the *ε* amino group. The very hydrophobic nature of this tunnel also explains why substitution of H3K79 with norleucine, which lacks the charged *ε* amino group but has the same 4-carbon aliphatic side chain, stabilizes the active complex.

### Conclusion and Implications

Our study reveals the basis for cross-talk between histone H2B ubiquitination and methylation of H3K79 by Dot1L and suggests a mechanism by which H2B-Ub increases the catalytic rate of Dot1L. By restricting the position of Dot1L on the face of the nucleosome, ubiquitin conjugated to H2B-K120 greatly reduces the orientations Dot1L can sample, while positioning the active site of Dot1L in an orientation that faces the nucleosome (Figure 1). The rigid nature of the Dot1L ubiquitin-binding element and its contacts with a region of ubiquitin that includes tail residue I73 (Figures 2b and S5d) help to limit Dot1L’s range of motion by partly immobilizing the flexible C-terminal ubiquitin tail. The range of motion is further limited by the contact between R282 and the nucleosome acidic patch (Figure 2d) which can occur in both the poised and active states. Conformational restriction of Dot1L thus increases its catalytic rate (McGinty et al., 2009) by greatly limiting the enzyme’s search for its substrate lysine. We speculate that the additional binding energy conferred by ubiquitin may further help pay the energetic cost of distorting histone H3 and lowering the pK_a_ of the substrate lysine. This energetic trade-off may also explain why H2B ubiquitination does not decrease the apparent K_M_ of the reaction, but instead affects k_cat_ (McGinty et al., 2009). Binding of the H4 tail to the opposing end of Dot1L further restricts movement of the N-terminal portion of Dot1L and position the active site over H3K79.

Conformational restriction could also account for the ability of H2B-Ub to stimulate methylation of H3K4, which lies in the long flexible N-terminal tail of histone H3 (Davey et al., 2002; Sun and Allis, 2002). The H3K4 methyltransferase belongs to the Set domain family, which is unrelated to the Dot1 family, and is part of the multiprotein yeast COMPASS and human SET1/MLL complexes (Miller et al., 2001). The ubiquitin L70-L73 patch required to stimulate Dot1L is also required to stimulate COMPASS (Holt et al., 2015), although mutations in the basic H4 tail residues that are required for Dot1 activity have no effect on H3K4 methylation (Fingerman et al., 2007). Structures of a 5- and 6-protein COMPASS subcomplex were recently reported (Hsu et al., 2018; Qu et al., 2018), but it is not known how this complex recognizes ubiquitin or the nucleosome. We speculate that contacts with ubiquitin anchor COMPASS to the nucleosome surface and reduce the conformational search for the flexible H3 tail.

The ability of Dot1L to reconfigure residues in the globular histone core raises the possibility that other histone-modifying enzymes may induce local conformational changes to access core residues that are fully or partly inaccessible. While histones can partially unfold when they are outside the histone octamer and bound to histone chaperones (English et al., 2006) or histone-modifying enzymes (Zhang et al., 2018), histones incorporated into nucleosomes adopt stable, virtually invariant conformations. However, recent methyl-TROSY NMR and cross-linking studies (Sinha et al., 2017) have suggested that the histone octamer can be distorted due to the action of ATP-dependent nucleosome remodeling enzymes. Our structural study of Dot1L has, to our knowledge, captured the first structural snapshot of a distortion in the globular histone core of an intact nucleosome. The view of the nucleosome core as a rigid substrate and binding partner should thus be modified to take into account structural variations, particularly in residues that lie at the ends of histone fold helices and in loops.

Because of its causative role in MLL Leukemia, Dot1L is actively pursued as a drug target (Chen and Armstrong, 2015). Most Dot1L inhibitors described to date are derived from S-adenosyl-L-homocysteine (SAH), a product of the methyltransferase reaction that binds to many other cellular enzymes, including other methyltransferases (Chen and Armstrong, 2015). The binding groove for the H4 tail revealed by our study suggests an approach to designing more specific inhibitors that bind to both the H4 tail binding region and the active site tunnel, thereby yielding Dot1L inhibitors that could serve as effective chemotherapeutic agents

## Author contributions

E.J.W. purified proteins and DNA, assembled nucleosomes, generated complexes, prepared EM grids, determined the structure of the poised state, and determined the structure of the active state in collaboration with N.H. C.H. cloned mutants of Dot1L. E.J.W., N.H. and C.W. wrote the manuscript, with input from C.H.

## Acknowledgements

We thank Xiangbin Zhang for purifying mutants, Duncan Sousa for advice. This research was supported, in part, by the National Cancer Institute’s National Cryo-EM Facility at the Frederick National Laboratory for Cancer Research under contract HSSN261200800001E. E.J.W. is a Damon Runyon Fellow supported by the Damon Runyon Cancer Research Foundation (DRG 2308-17), N.H. is supported by an EMBO Long-term Fellowship (EMBO-ALTF 309-2017), and a grant from the National Institute of General Medical Sciences (GM095822).

## Data deposition

The cryo-EM maps described in this study have been deposited in the EMDB with accession codes EMD-XXX (2.96 Å active Dot1L-nucleosome) and EMD-YYY (3.9 Å poised Dot1L-nucleosome). Coordinates of the corresponding atomic models were deposited in the Protein Data Bank under accession code XXX (active Dot1L-nucleosome) and YYY (poised Dot1L-nucleosome).

---

**Supplementary Table 1:** Data collection and refinement statistics.

Cryo-EM data collection, refinement and validation statistics
|  | Active State<br>(EMDB-xxxx)<br>(PDB xxxx) | Poised State<br>(EMDB-xxxx)<br>(PDB xxxx) |
| --- | --- | --- |
| **Data collection and processing** |  |  |
| Magnification | 130,000 | 130,000 |
| Voltage (kV) | 300 | 300 |
| Electron exposure (e-/Å<sup>2</sup>) | 50 | 50 |
| Defocus range (μm) | -1 to -2.5 | -1 to -2.5 |
| Pixel size (Å) | 0.532 | 0.532 |
| Symmetry imposed | C2 |  |
| Initial particle images (no.) | 800727 | 868027 |
| Final particle images (no.) | 23778 | 108658 |
| Map resolution (Å) | 2.96 | 3.9 |
| FSC threshold | (0.143) | (0.143) |
| Map resolution range (Å) | 2.96-272.4 | 3.9-272.4 |
| **Refinement** |  |  |
| Initial model used (PDB code) | 1KX3, 1NW3,<br>1UBQ | 1KX3, 3QOW<br>1UBQ |
| Model resolution (Å) | 3.2 | 4.40 |
| FSC threshold | (0.5) | (0.5) |
| Model resolution range (Å) | 272.4-3.3 | 272.4-4.4 |
| Map sharpening *B* factor (Å<sup>2</sup>) | -73 | -180 |
| Model composition |  |  |
| Non-hydrogen atoms | 15510 | 14957 |
| Protein residues | 1186 | 1126 |
| Nucleotides | 292 | 292 |
| Ligands | 1 | 0 |
| *B* factors (Å<sup>2</sup>) |  |  |
| Protein | 71.6 | 113.11 |
| Nucleotide | 19.47 | 134.41 |
| Ligand | 77.42 | - |
| R.m.s. deviations |  |  |
| Bond lengths (Å) | 0.013 | 0.009 |
| Bond angles (°) | 1.04 | 1.35 |
| Validation |  |  |
| MolProbity score | 1.46 | 1.41 |
| Clashscore | 2.72 | 4.29 |
| Poor rotamers (%) | 0.99 | 0.94 |
| Ramachandran plot |  |  |
| Favored (%) | 94.08 | 96.73 |
| Allowed (%) | 5.92 | 3.27 |
| Disallowed (%) | 0 | 0 |

**Supplementary Table 2:** Previous mutants and their effects on H3K79 methylation by Dot1L. Previous mutations that probe the activity of Dot1L are listed.

| Target | Mutation | Organism | H3K79 methylation | *In vivo/in vitro* | Citation |
| --- | --- | --- | --- | --- | --- |
| H2B-Ub | H2B K123R | Yeast | abolished | *In vivo* | (Briggs et al., 2002; Ng et al., 2002) |
| | ΔRad6 | Yeast | abolished | *in vivo* | (Briggs et al., 2002; Ng et al., 2002) |
|  | Rnf20 (knockdown) | Human | reduced | *in vivo* | (Wang et al., 2013) |
|  | Rnf20 (overexpression) | Human | increased | *in vivo* | (Zhu et al., 2005) |
| Ubiquitin | L71A, L73A | Human | decreased | *in vitro* | (Holt et al., 2015) |
|  | L8,I44A | Human | no effect | *in vitro* | (McGinty et al., 2009) |
| Acidic patch | H2A E64A | Human | no effect | *in vitro* | (McGinty et al., 2009) |
|  | H2A E68A | Human | no effect | *in vitro* | (McGinty et al., 2009) |
| H4 tail | R19A, R17A | Human | abolished | *in vitro* | (McGinty et al., 2009) |
|  | R19A, R17A | Yeast | abolished | *in vivo/in vitro* | (Altaf et al., 2007; Fingerman et al., 2007) |
|  | R19E, R17E | Yeast | abolished | *in vivo* | (Fingerman et al., 2007) |
| | $\Delta$ 17-20 | Yeast | abolished | *in vivo/in vitro* | (Fingerman et al., 2007) |
| Dot1L | E374 | Yeast | abolished | *in vitro* | (Sawada et al., 2004) |
|  | Y350 | Yeast | abolished | *in vitro* | (Sawada et al., 2004) |
|  | W543A | Yeast | abolished | *in vitro* | (Sawada et al., 2004) |
|  | W543F | Yeast | abolished | *in vitro* | (Sawada et al., 2004) |

**Supplementary Figure S1:**
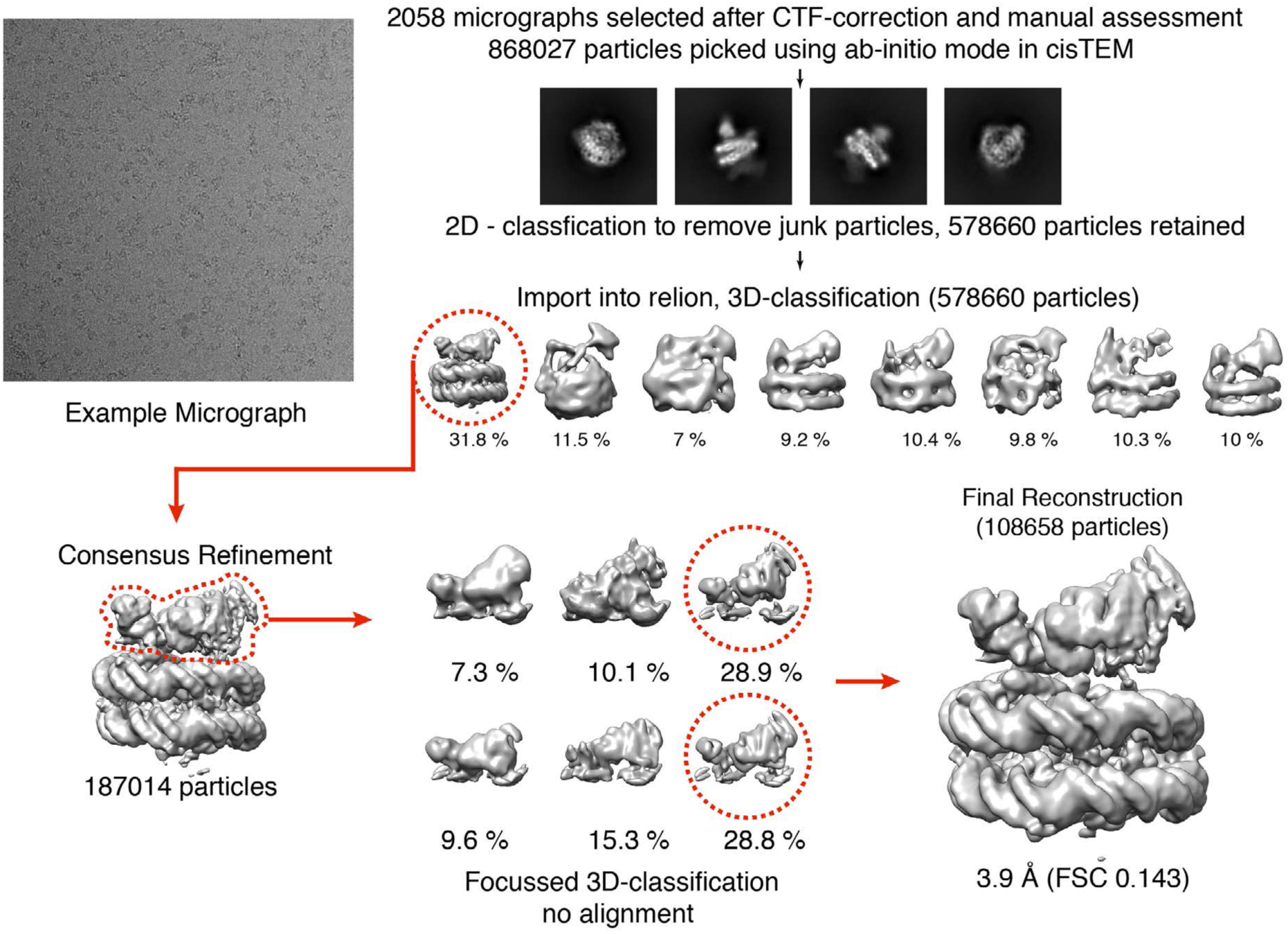
Cryo-EM processing pipeline for the poised Dot1L-nucleosome structure. The displayed micrograph is low-pass filtered to 20Å for improved particle visibility.

**Supplementary Figure S2:**
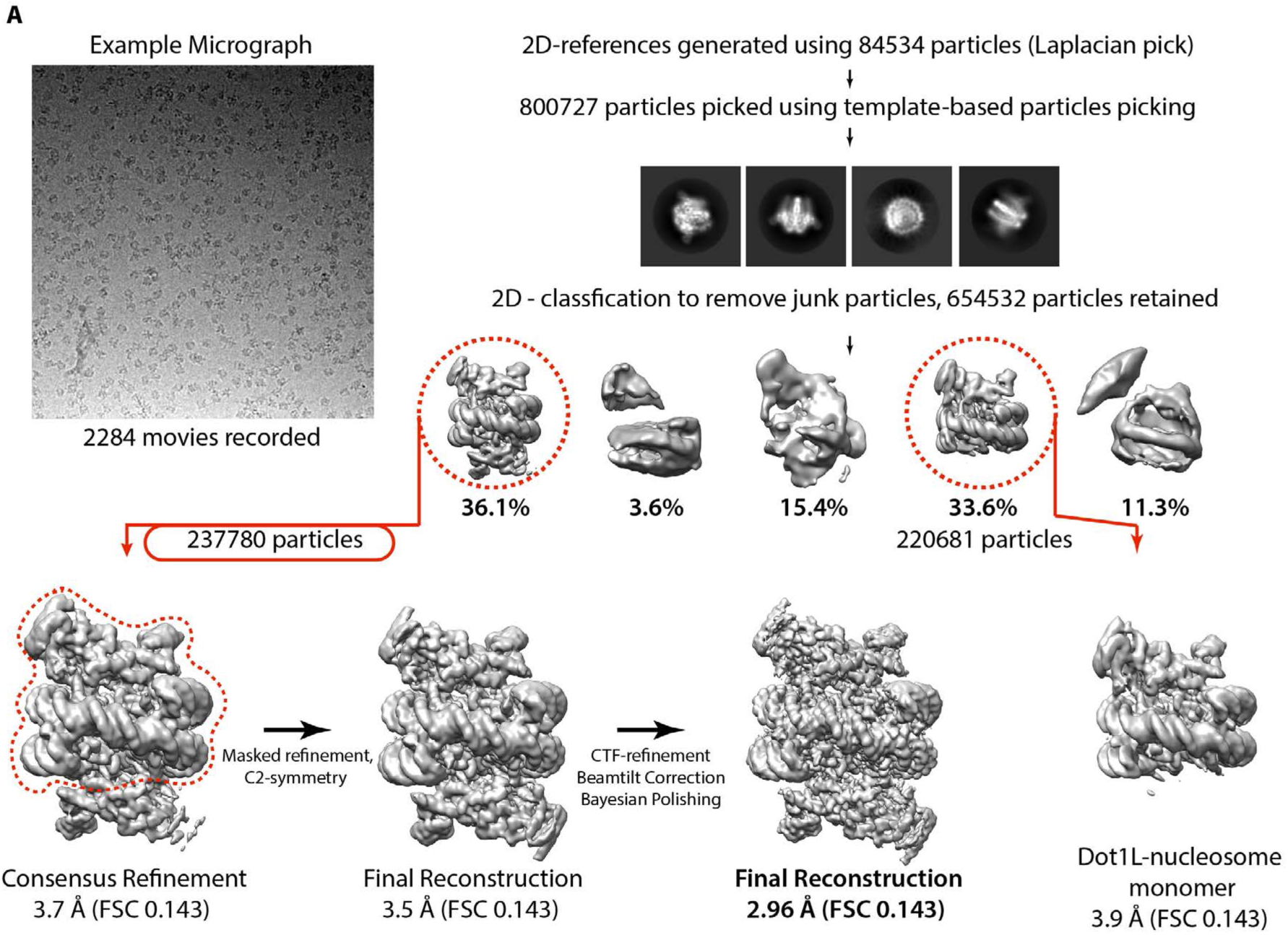
Cryo-EM processing pipeline for the active Dot1L-nucleosome structure. The displayed micrograph is low-pass filtered to 20Å for improved particle visibility. The resolution improvement from 3.5Å to 2.96Å was achieved by Relion 3’s particle-based CTF-refinement, beamtilt correction and bayesian polishing tools. These features were applied in an iterative manner as listed until no resolution and map improvement could be achieved (this corresponded to 2 cycles).

**Supplementary Figure S3:**
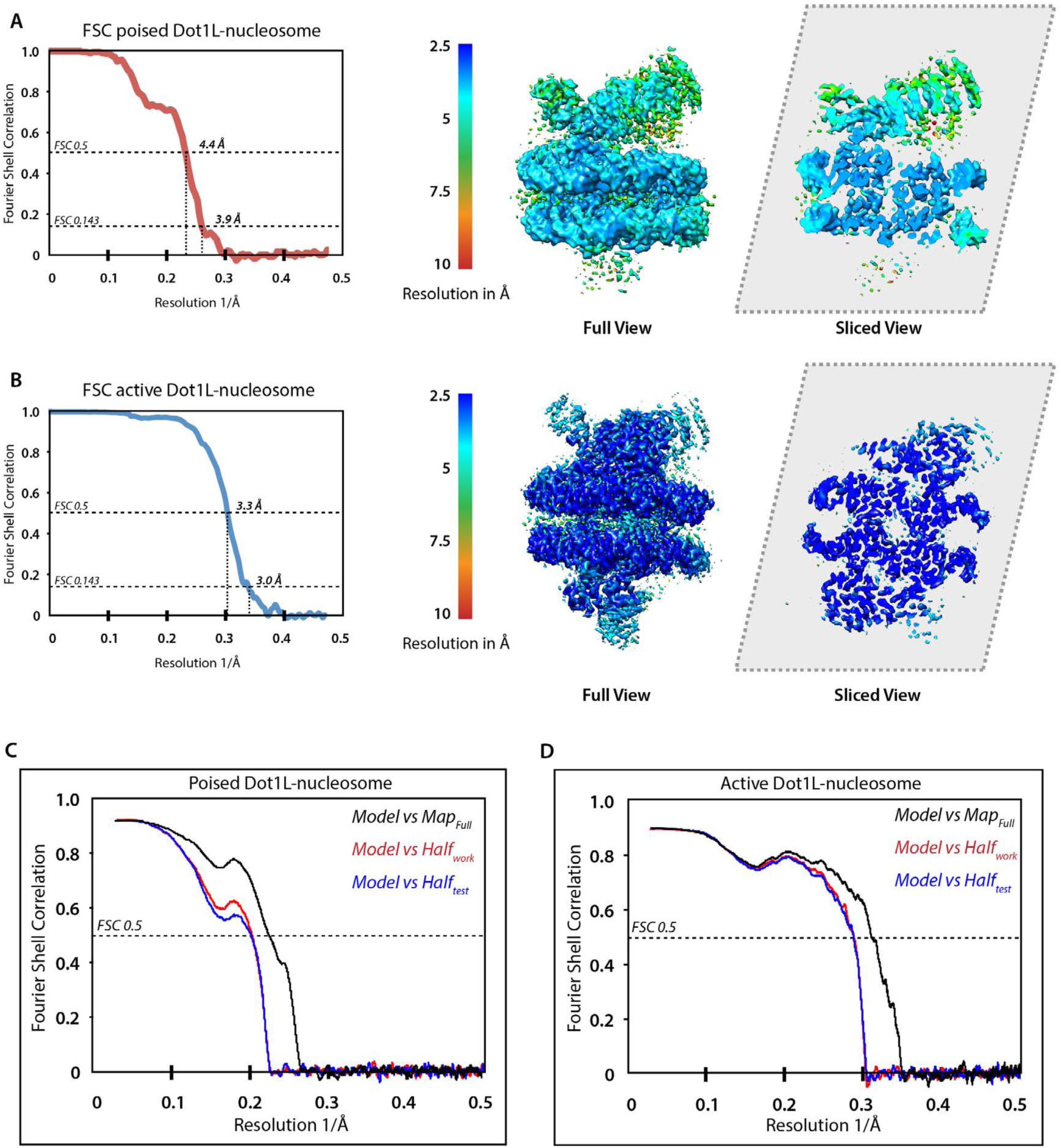
Cryo-EM map and model validation. **a**, Fourier Shell Correlation (FSC) and local resolution assessment by Resmap for the poised Dot1L-nucleosome reconstruction. The final resolution as determined by FSC 0.143 criterion is 3.9Å. The local resolution in Å is displayed on the sharpened full map (middle = surface, right = cross section) of poised Dot1L. **b**, FSC and local resolution assessment for the active Dot1L-nucleosome structure. The final resolution is 2.96Å as determined by the FSC 0.143 criterion. The local resolution in Å is displayed on the sharpened full map of active Dot1L, similar to B. **c**, FSC curves calculated between the refined atomic model of poised Dot1L-nucleosome complex and the masked, sharpened half map used for the refinement are shown in red (FSC_work_), and of the refined atomic model against the second masked, sharpened half-map in blue (FSC_test_). We also plotted the FSC of the refined model against the full map displayed in black (FSC_Full_). The close agreement between the FSC_work_ and FSC_test_ curves and the absence of significant correlation beyond the calculated map resolution for all three curves indicates absence of overfitting and model bias. **d**, Same as c, but for active Dot1L.

**Supplementary Figure S4:**
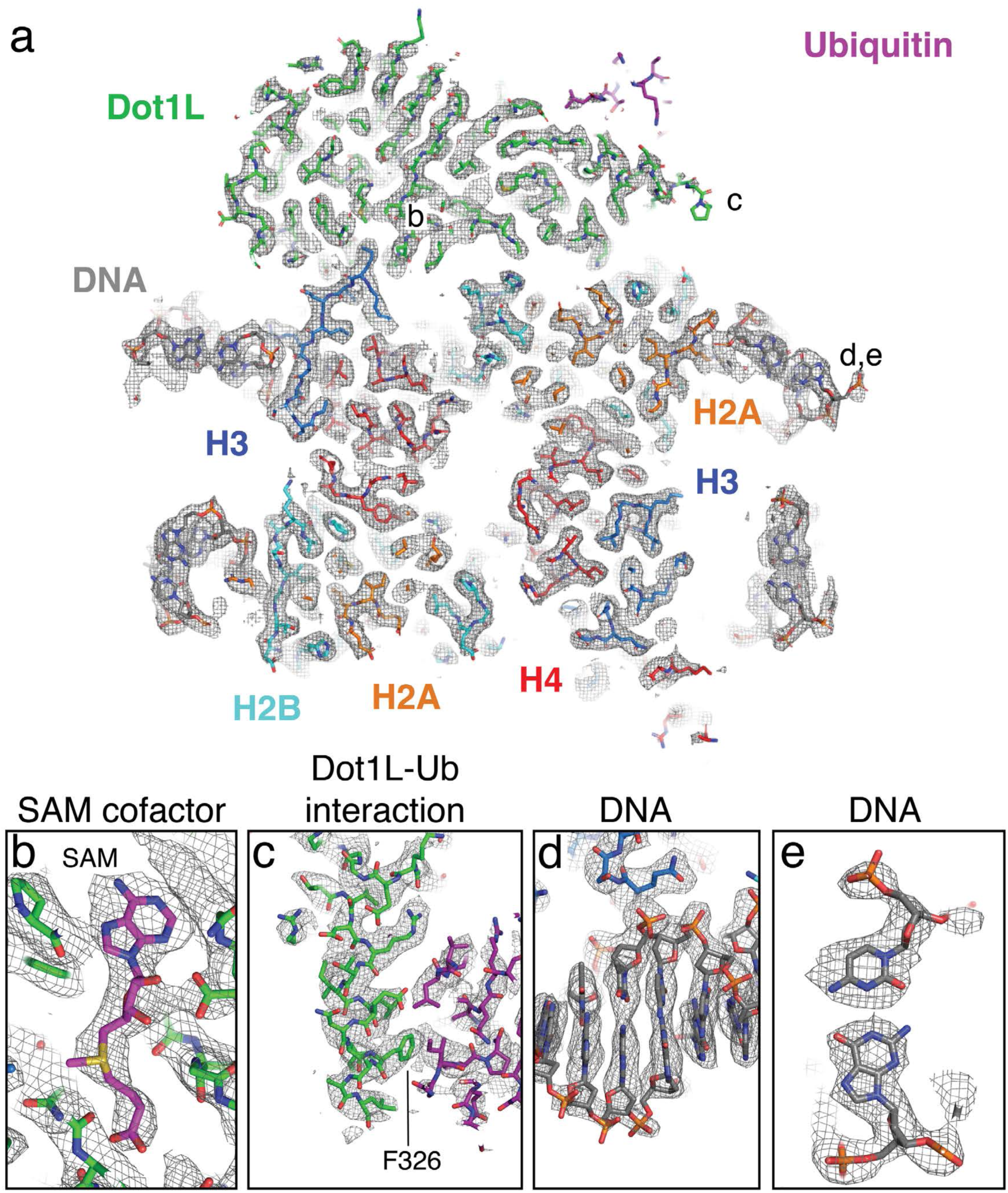
Example density of the active-state EM structure. **a**, A vertical slice through the active state structure centered on H3K79. Atomic models of all chains are shown as sticks and the EM density is shown as a gray mesh. Areas highlighted in the boxes are denoted with letters b, c, d and e. **b** Example density of the SAM cofactor. **c**, Example density of the interaction between ubiquitin and the Dot1L C-terminal helix. **d.e**, Example EM density of the DNA.

**Supplementary Figure S5:**
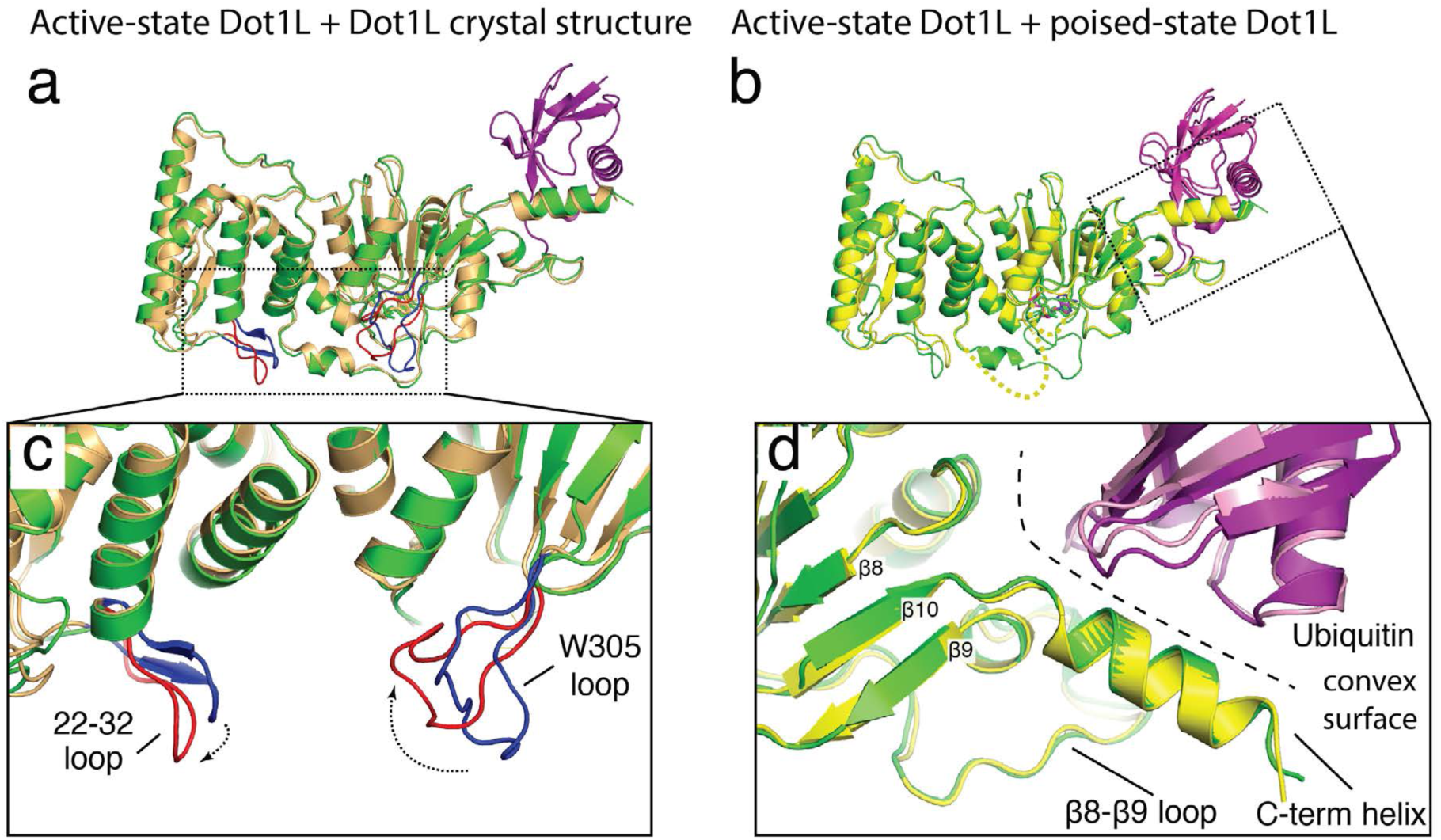
Comparisons between Dot1L in different states. **a** Superimposition between the active state Dot1L and the Dot1L crystal structure (PDB ID: 1NW3). **b**, Superimposition between the active state and poised sate Dot1L. The disordered Dot1L F131 and W305 loops in the poised state are depicted as dashed yellow lines. **c**, Close up view of loop restructuring that occurs in Dot1L during the transition to the active state. The active state loops are colored red and the loops from the crystal structure of Dot1L (PDB ID: 1NW3) are colored blue. **d**, Close up view of the superimposed Dot1L-ubiquitin interaction in the poised and active states showing that the contact surface does not change. The convex surface made up by the Dot1L C-terminal helix and the *β*8- *β*9 loop is depicted as a dashed line.

**Supplementary Figure S6:**
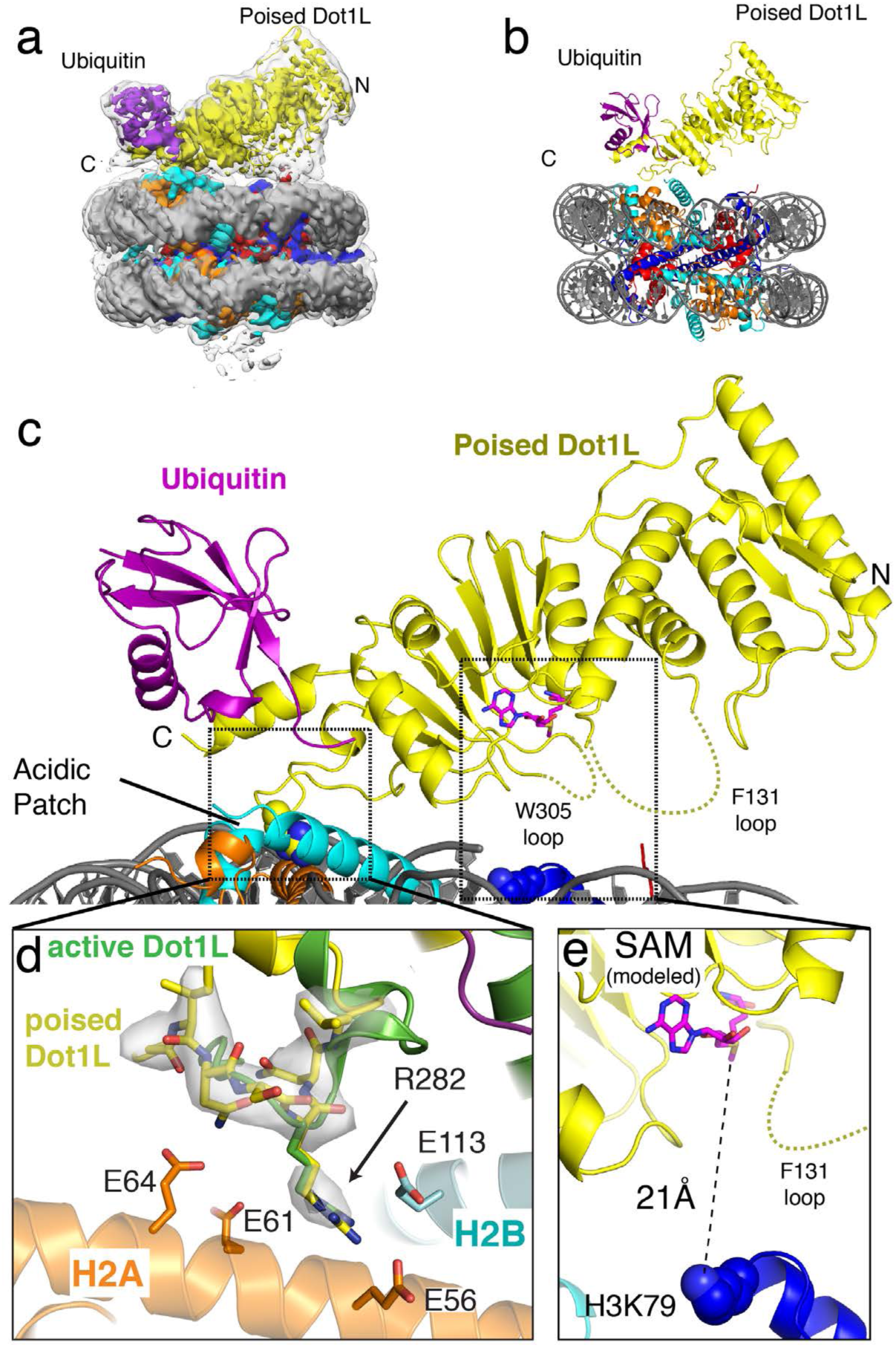
Details of the poised state structure. **a**, EM density for the poised-state structure. The unsharpened density is shown as a transparent surface and the sharpened density is shown as an opaque surface colored as in the main text. The N- and C-terminal parts of Dot1L are denoted with C and N. **b**, The poised state structure atomic model colored as in the main text. **c**, Close up view of Dot1L in the poised state showing that the only direct contact between Dot1L and the nucleosome is through the acidic patch. The F131 and W305 loops do not have defined density in the poised state and are depicted as yellow dashed lines. **d**, Detailed view of the poised state interaction with the acidic patch superimposed with the active state structure. The poised state structure is colored yellow and the active state structure is colored green. Sharpened EM density for the poised state structure is shown as a transparent gray surface and acidic patch residues of H2A and H2B are shown as sticks. **e**, Close up view of the poised state Dot1L active site showing that a modeled SAM cofactor is too far for the methyl transfer reaction to occur.

**Supplementary Figure S7:**
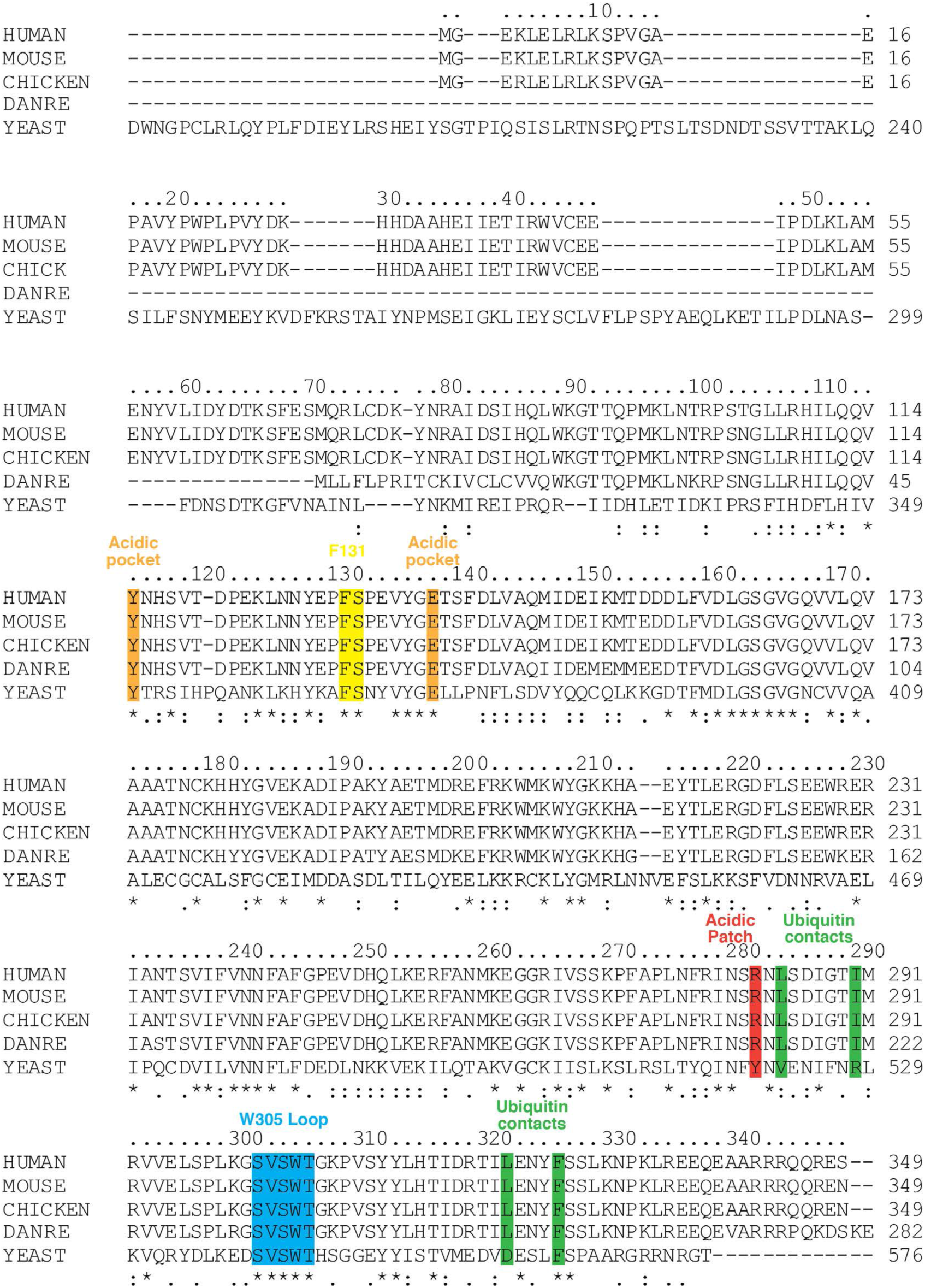
Dot1L conservation across species. Multiple Sequence alignment of Dot1L variants from different species. *Homo sapiens (Human*, Q8TEK3), *Mus musculus (Mouse*, Q6XZL8), *Gallus gallus* (Chicken), *Danio rerio (Zebrafish*, F1Q4W7), *Saccharomyces cerevisiae* (Yeast, Q04089). The alignment was performed with the Clustal Omega program.

**Supplementary Figure S8:**
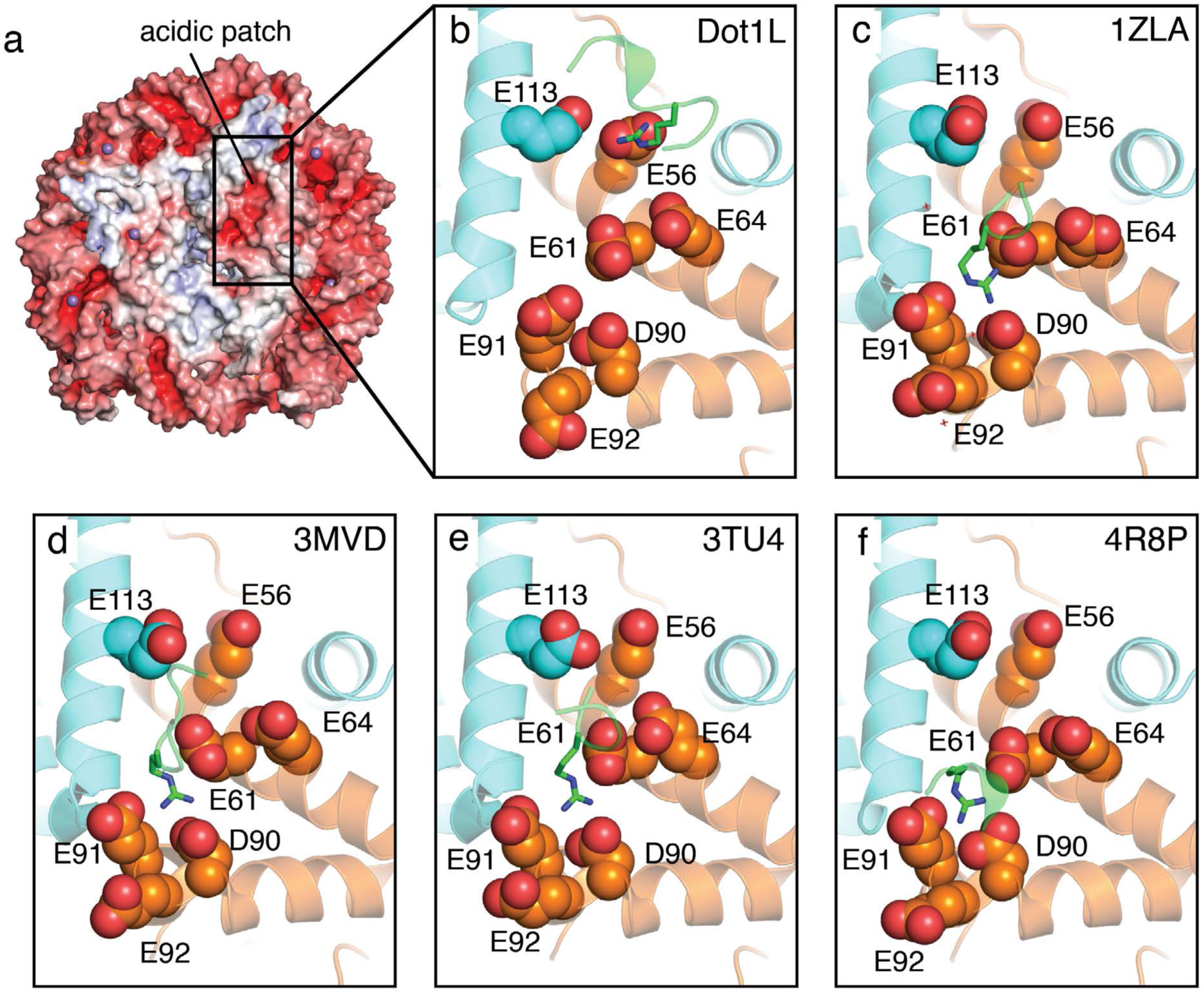
Arginine anchor interactions with the acidic patch. **a**, Surface representation of the nucleosome colored by electrostatic potential. The highly charged acidic patch is indicated by the black square. **b-f**, Close up views of the arginine anchor interaction with the acidic patch for different protein-nucleosome complexes. **b**, Dot1L, **c**, LANA peptide (PDB: 1ZLA), **d**, RCC1 (PDB: 3MVD), **e**, Sir3 (3TU4), **f**, PRC1 Ubiquitylation Module (4R8P).

**Supplementary Figure S9:**
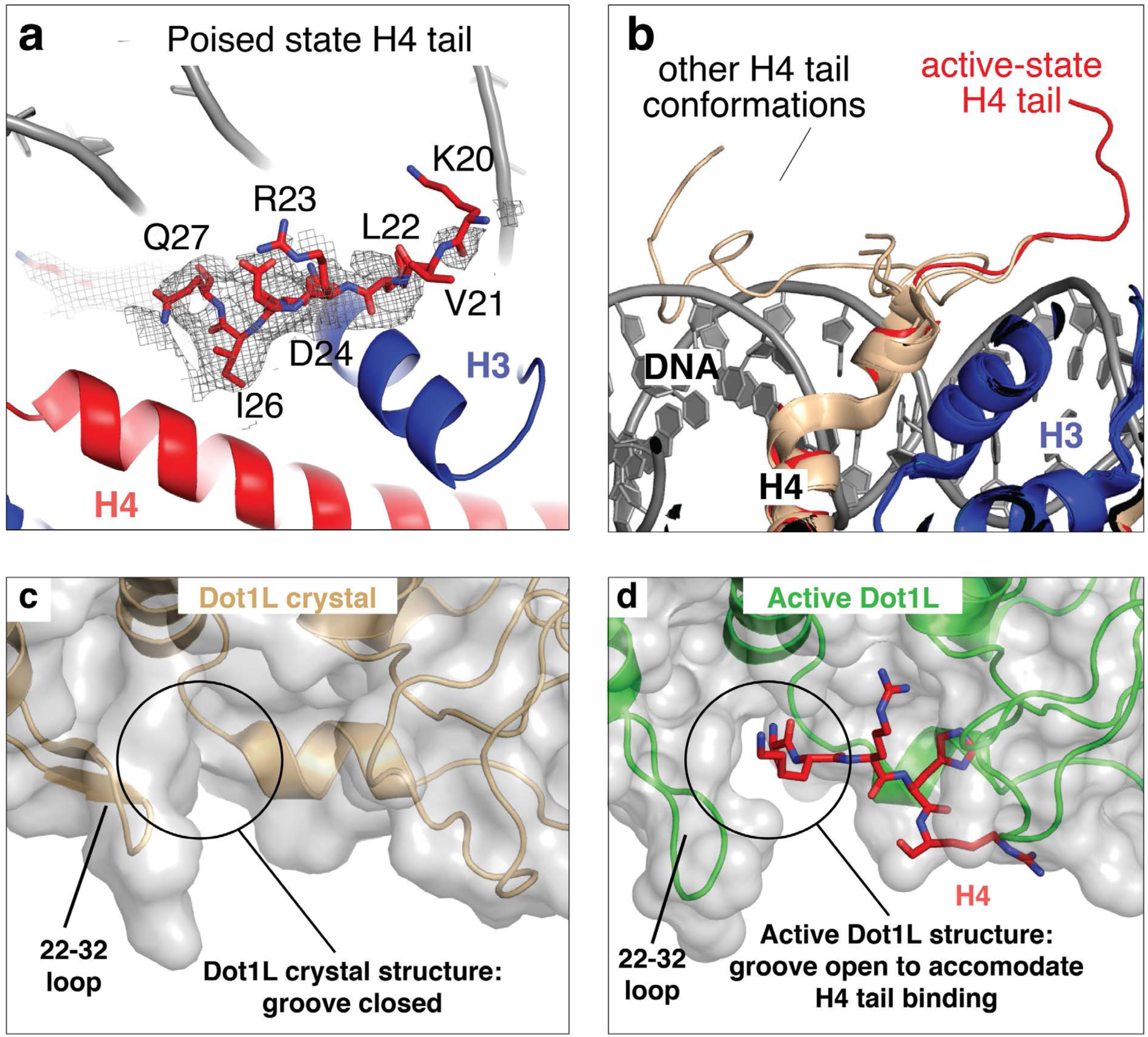
Details of the H4 tail interaction with Dot1L. **a**, Example density of the H4 tail in the poised state structure is depicted as a gray mesh and shows that residues after K20 are not structured. **b** The active state Dot1L structure (red) is superimposed with several high resolution nucleosome crystal structures (PDB ID: 1KX3, 1KX5, 5YOD, 1S32, 3UTA, 5Y0C, 3C1B, 3UT9 and 3UTB) colored tan showing that the H4 tail is usually unstructured or associated with DNA **c**, The Dot1L crystal structure (PDB ID: 1NW3) is depicted in cartoon representation and colored brown with a semi-transparent gray surface. The groove for H4 tail is closed in this state. **d**, The active Dot1L structure is depicted in cartoon representation and colored green with a semi-transparent gray surface. The groove for the H4 tail is open in this state.

**Supplementary Figure S10:**
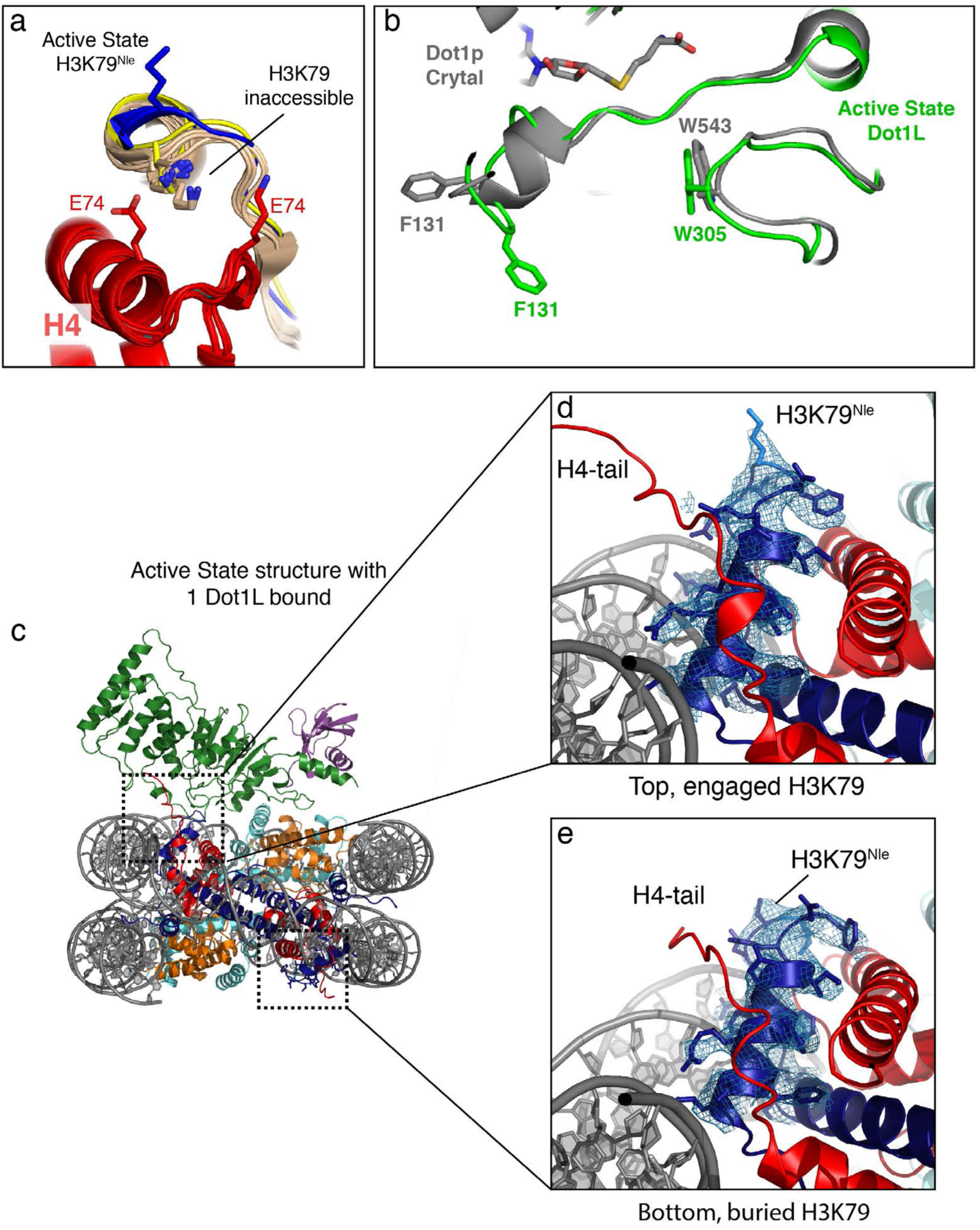
Details of the H3K79 conformational change. **a**, The active state Dot1L structure (blue) is superimposed with the poised Dot1L structure (yellow) and several high resolution nucleosome crystal structures (PDB ID: 1KX3,1KX5, 5YOD, 1S32, 3UTA, 5Y0C, 3C1B, 3UT9 and 3UTB) colored tan showing that the sidechain of H3K79 is always held in an inaccessible state. **b**, Superimposition of the active state Dot1L (green) the yeast Dot1p crystal structure (PDB ID: 1U2Z, gray) showing the relative conformations of the W305 and F131 loops. The yeast Dot1p residue W543 is in a similar position as W305 in Dot1L. **c**, Overview of the active state structure where only one Dot1L is present. **d**, Close up view of the H3K79^Nle^ loop on the side of the nucleosome that is bound by Dot1L showing that H3K79^Nle^ is oriented upward. **e**, Close up view of the H3K79^Nle^ loop on the side of the nucleosome that is not bound by Dot1L, showing that H3K79^Nle^ is oriented in the inaccessible conformation.

## Methods

### Protein and DNA preparations

#### Purification of Dot1L variants

A plasmid bearing the gene for human Dot1L, MSCB-hDot1Lwt, was a gift from Yi Zhang (Addgene plasmid # 74173; http://n2t.net/addgene:74173; RRID: Addgene_74173). A truncated Dot1L, Dot1L(2-416), was amplified by PCR from MSCB-hDot1Lwt using primers that encoded KpnI and EcoRI restriction sites and a TEV protease site. The resulting PRC product was cloned into pET32-a (Novagen) using the KpnI and EcoRI restriction sites to produce the expression constructs, which contained an N-terminal fusion of thioredoxin (Trx), a hexahistidine tag and a cleavage site for Tobacco Etch Virus (TEV) protease (Trx-His-TEV-Dot1L(2-416)). Dot1L point mutants were cloned in the Trx-His-TEV-Dot1L(2-416) plasmid using around-the-horn mutagenesis. Rosetta2(DE3) cells were transformed with the Trx-His-TEV-Dot1L(2-416) plasmid and grown in 2XYT media at 37°C while shaking. When the cell density reached an OD_600_ of 0.8-1.0, the growth temperature was reduced to 18°C, 1mM IPTG was added to induce protein expression, and the cells were grown overnight. Cells were harvested by centrifugation and the cell pellet was resuspended in 20 ml of Dot1L lysis buffer (50 mM Tris pH 8.0, 750 mM NaCl, 5% glycerol, 0.5 mM dithiothreitol (DTT), 15 mM imidazole, 100 μM Phenylmethylsulfonyl fluoride (PMSF) per liter of growth medium, flash frozen in liquid nitrogen, and stored at −80°C until needed.

To purify Dot1L, the frozen pellet was thawed at 25°C and cooled on ice. The cells were lysed using an LM10 Microfluidizer (Microfluidics) and the clarified lysate was added to 10 ml of Ni-NTA beads (Qiagen) that had been equilibrated in NiA buffer (50 mM Tris pH 8.0, 750 mM NaCl, 5% glycerol, 0.5 mM DTT, 15 mM imidazole). The beads were incubated with the lysate at 4°C for 45 minutes, poured into a gravity flow column and washed with 200 ml of NiA buffer. Protein was eluted using ~ 70ml of NiB buffer (50 mM Tris pH 8.0, 750 mM NaCl, 5% glycerol, 0.5 mM DTT, 250 mM imidazole). After elution the absorbance at 280nm was measured and 6mg of TEV protease was added directly to the eluate and dialyzed overnight at 4°C against Dot1L Dialysis buffer (50 mM HEPES, pH 7.0, 100 mM NaCl, 5% glycerol, 1 mM DTT). Precipitate was removed by centrifugation and filtration through a 0.22μm Millex syringe filter (Millipore). To separate Dot1L from TEV and the cleaved Trx-His tag, the sample was loaded onto a 5 ml SP HP column (GE Healthcare) cation exchange column equilibrated in ion exchange buffer (50 mM HEPES, pH 7.0, 50 mM NaCl, 5% Glycerol, 1 mM DTT) and eluted with a 50 mM − 1000 mM linear NaCl gradient of 15 column volumes. Peak fractions corresponding to Dot1L were pooled and the buffer exchanged into Dot1L storage buffer (30 mM Tris pH 8.0, 200 mM NaCl, 1 mM Tris(2-carboxyethyl)phosphine (TCEP), 20% glycerol) by repeated concentration and dilution using an Amicon Ultra spin concentrator (Millipore). The pure concentrated protein (~500μM) was flash frozen and stored at −80°C until use.

Dot1L(2-316) was purified in the same manner as Dot1L(2-416) except that, after TEV cleavage, the protein was concentrated using an Amicon Ultra spin concentrator (Millipore) and separated from TEV and the Trx-His tag using a Superdex 200 16/600 size exclusion chromatography column (GE Healthcare) that was pre-equilibrated with SEC buffer (30 mM Tris pH 8.0, 200 mM NaCl, 2 mM DTT, 5% glycerol). Peak fractions were pooled, the buffer exchanged into Dot1L storage buffer and frozen as described for Dot1L(2-416).

#### Purification of His-TEV-ubiquitin(G76C)

His-TEV-Ubiquitin(G76C) was purified as described previously(Morgan et al., 2016). Briefly, His-TEV-ubiquitin was expressed in Rosetta2 cells and purified by Ni-affinity chromatography, followed by dialysis into ammonium acetate, pH 4.5, overnight. Dialyzed protein was loaded onto tandem SP-HP cation exchange columns (GE) that were pre-equilibrated with ammonium acetate pH 4.5. Ubiquitin was eluted using a step gradient of 750mM NaCl, dialyzed into Ubiquitin storage buffer (10 mM Tris pH 7.8, 1 mM TCEP), flash frozen and stored at −80°C.

#### Purification of histone proteins

Unmodified *Xenopus laevis* histone expression plasmids containing H2A, H2B, H3 and H4 were a generous gift from Greg Bowman. Histone expression and purification were conducted following standard protocols described in Luger et. al. (Dyer et al., 2004).

To produce histone H3 with the unnatural amino acid, norleucine, in place of K79, histone H3 was cloned into pQE-81L and K79 was mutated to methionine around-the-horn PCR. For protein expression, the methionine auxotroph, *E. coli* B834(DE3)pLysS (Novagen) was transformed with the pQE-81L(H3K79M) plasmid and grown at 37°C in M9 minimal media supplemented with 60 mg/L of all amino acids except for 100 mg/L of lysine and 500 mg/L of methionine. Cells were grown to an OD_600_ of 0.6, pelleted by centrifugation and then washed three times with M9 medium. The cells were resuspended in fresh M9 media supplemented with all amino acids except methionine and supplemented with 500 mg/L of norleucine. 1mM IPTG was added to induce expression and the cells were grown at 37°C for 2-3 hours. Cells were harvested by centrifugation and the H3K79Nle histone was purified as described in Luger et. al.(Dyer et al., 2004)

#### Purification of 601 DNA

The Widom 601 strong positioning DNA sequence(Lowary and Widom, 1998) was produced in the E.coli strain XL-1 Blue using the pST55-16x601 plasmid(Makde et al., 2010). The pST55-16x601 plasmid was purified and 601 DNA sequence was isolated from the plasmid as described previously (Dyer et al., 2004).

#### Preparation of H2B-Ubiquitin

H2B-Ub containing a dichloroacetone linkage between the ubiquitin C-terminus and H2BK120 was prepared as described(Morgan et al., 2016). Briefly, 100uM His-TEV-Ubiquitin(G76C) and 100uM H2B(K120C) were mixed in crosslinking buffer (50 mM Borate pH 8.1, 1 TCEP) and incubated at 50°C for one hour to reduce cysteines. The mixture was cooled on ice and then crosslinking was initiated by the addition of 100uM dichloroacetone and allowed to proceed for 1 hour. The reaction was then quenched with 50mM β-mercaptoethanol (BME), dialyzed into water and lyophilized. The product was then resuspended in NiNTA-6MU buffer (6M urea, 50mM Tris pH7.8, 500mM NaCl, 10mM BME, 20mM Imidazole, 0.1mM PMSF) and incubated with 20 ml of Ni-NTA resin. Unreacted H2B and H2B-H2B crosslinks were removed in the flow through. The desired H2B-Ub product along with unreacted Ubiquitin and Ubiquitin-Ubiquitin crosslinks were eluted in NiNTA-6MU buffer supplemented with 250mM Imidazole and dialyzed into TEV cleavage buffer (50 mM Tris-HCl pH 8.0, 0.5 mM EDTA, and 1 mM DTT) and the his-tag on ubiquitin was removed by the addition of TEV protease. The material was dialyzed into water, lyophilized, re-dissolved in NiNTA-6MU and uncleaved protein and TEV were removed by passage over Ni-NTA resin. The flow through was dialyzed into water and lyophilized again. Finally, the dry protein was dissolved in 7M Guanidine-HCl and separated on a Proto 300 C4 column (Higgins Analytical) in 0.1% TFA and a gradient of 0.1%TFA + 90% acetonitrile. Fractions corresponding to pure H2B-Ub were pooled and lyophilized.

#### Reconstitution of nucleosome core particles

Nucleosome core particles containing either unmodified, wild type H2B, H2B-Ub, or norleucine-substituted H2B-Ub were reconstituted as described previously(Dyer et al., 2004; Luger et al., 1999). Briefly, histone octamers were prepared from individual histone proteins by refolding and salt dialysis. Histone octamers were isolated from H2A/H2B dimers and H3/H4 tetramers by size exclusion chromatography, concentrated using an Amicon Ultra spin concentrator (Millipore) and stored at 4°C until use. Nucleosomes were prepared by mixing histone octamers and purified 601 DNA under high salt followed by salt gradient deposition over 18 hours. Fully formed nucleosomes were separated from misfolded nucleosomes by separation on TSK DEAE-5PW HPLC column(Luger et al., 1999) (TOSOH biosciences).

### Dot1L activity assays

#### Endpoint methylation assays

Stocks of 100 nM of each mutant Dot1L(2-416) plus 40 μM S-adenosyl methionine SAM and 2 μM H2B-Ub nucleosome or unmodified wild type nucleosome were prepared in assay buffer (20mM Tris pH 8.0, 50mM NaCl, 1mM DTT, 0.1mg/ml bovine serum albumin). The reactions were initiated by mixing 6 μl of Dot1L and 6μl of nucleosome for a final concentration of 50 nM Dot1L and 1μM nucleosome. The reaction proceeded for 30 min at 25°C and was quenched by addition of 3μl 0.5% TFA. 10 μl of the quenched reaction was transferred to a white 384 well plate and S-adenosyl homocysteine (SAH) production was detected using the MTase Glo assay (Promega). Each reaction was done in triplicate and the standard deviation of the raw data is reported.

### Cryo-EM sample preparation and structure determination

#### Poised state Dot1L

A 4:1 complex between Dot1L(2-416) (60 μM) and H2B-Ub nucleosome (15 μM) was prepared in crosslinking buffer (50 mM HEPES, pH8.0, 50 mM NaCl, 1mM EDTA, 0.2 mM DTT) in the presence of 1 mM 5’-(diaminobutyric acid)-N-iodoethyl-5’-deoxyadenosine ammonium hydrochloride (AAI), an N-mustard SAM analog inhibitor(Weller and Rajski, 2006) that was a generous gift from Dr. Lindsay Comstock-Ferguson (Wake Forest). The mixture was incubated for 1 hour at 30°C and then diluted to 100 nM nucleosome in crosslinking buffer. Glutaraldehyde crosslinking was initiated by adding an equal volume of 0.06% glutaraldehyde to the mixture to yield a reaction containing 50 nM H2B-Ub nucleosome, 200 nM Dot1L and 0.03% glutaraldehyde. The reaction was incubated for 1 hour on ice and then quenched by addition of 100 mM Tris, then dialyzed overnight against quenching buffer (50mM Tris pH 8.0, 50mM NaCl, 1mM DTT) which removed the AAI inhibitor. The complex was then concentrated using an Amicon Ultra spin concentrator (Millipore) and purified on a Superdex 200 16-60 size exclusion column (GE Healthcare) that was pre-equilibrated with EM buffer (20 mM HEPES pH 7.5, 50 mM NaCl, 1 mM DTT). Peak fractions were concentrated to a final nucleosome concentration of 3.4 μM nucleosome. A sample at 1 mg/ml (2.43 μM) was frozen on 400 mesh C-Flat grids (Electron Microscopy Sciences) with 2 μm holes using a Vitrobot (Thermo Fisher Scientific) with settings at 4°C, 100% humidity and 3 sec blot time.

#### Active state Dot1L

A 4:1 complex containing 60 μM Dot1L(2-416) 15 μM H2B-Ub nucleosomes containing the H3K79Nle mutation was prepared in crosslinking buffer supplemented with 200 μM SAM. The resulting complex was diluted to 100nM using reaction buffer and 2 mM SAM was added to a final concentration of 160uM. An equal volume of 0.064% glutaraldehyde in reaction buffer was added to the complex to yield a crosslinking reaction containing 50 nM H2B-Ub nucleosome, 200 nM Dot1L and 0.032% glutaraldehyde. After 1 hour on ice, the reaction was quenched with 100 mM Tris and dialyzed against quenching buffer. The crosslinked complex was purified and frozen as described for the poised complex except that the concentration of the active complex on the grid was 0.75 mg/ml.

#### Refinement and model building

All data were collected at the National Cryo-Electron Microscopy Facility (NCEF) at the National Cancer Institute on a Titan Krios at 300 kV utilizing a K2 direct electron detector in Super resolution electron counting-mode at a nominal magnification of 130,000 and a pixel size of 0.532 Å. For both the poised and active state Dot1L datasets, data were collected at a nominal dose of 50 e^−^/Å^2^ with 40 frames per movie and 1.25 e^−^/frame. A total of 2,143 super resolution movies were collected for the poised state dataset and 2,284 movies were collected for the active state dataset.

For the poised Dot1L dataset, all movies were motion-corrected, dose-weighted and binned by a factor of 2 using MotionCorr2 (Zheng et al., 2017) which resulted in a pixel size of 1.064 Å. Contrast transfer function (CTF) correction was performed with Ctffind4 (Rohou and Grigorieff, 2015). Particle picking utilizing the ab-initio particle picking mode was performed in cisTEM (Grant et al., 2018), which resulted in 868,027 particles that were subjected to 2D classification to remove junk particles. A cleaned dataset of 578,660 particles was then transferred to Relion 2.0 (Kimanius et al., 2016) for 3D classification, yielding a single class with 187,014 particles that showed density for a well-resolved Dot1L in the poised state after refinement (Extended Data Figure 1). Focused classification without alignment using a mask that encompassed the Dot1L-ubiquitin complex further reduced the dataset to 108,658 particles, which were refined to 3.9 Å resolution as given by the Fourier shell correlation (FSC) 0.143 criterion (Rosenthal and Henderson, 2003; van Heel and Schatz, 2005) (4.4 Å at FSC 0.5). The final map was sharpened with an automatically calculated B-factor of −180 Å^2^ using the Relion post-processing tool.

The active Dot1L dataset was processed in Relion 3 (Zivanov et al., 2018). All movies were motion-corrected, dose-weighted and binned by a factor of 2 using the Relion 3 implementation of MotionCorr2 (Zivanov et al., 2018), which resulted in a pixel size of 1.064 Å. A initial batch of 143 micrographs were selected based on CTF fit statistics to pick 84,534 particles using the Laplacian auto-picking feature. Initial class averages were calculated and used to perform template-based particle picking on the entire dataset, yielding 800,727 particles. Next, 2D-classification was performed to remove junk particles, and 652,532 particles were retained and 3D-classified (Extended Data Figure 2). The best resolving, highest-populated class corresponded to a 2:1 Dot1L-nucleosome complex and contained 237,780 particles. A masked refinement including the nucleosome and the upper, better resolved Dot1L and imposed C2 symmetry yielded the a 3.5 Å reconstruction. The 2:1 Dot1L complex particle stack was subjected to 2 iterations of CTF refinement, beam tilt correction and Bayesian polishing as implemented in Relion 3.0. Refinement of the shiny particles using C2 symmetry and a mask that encompassed the nucleosome and the upper Dot1L and yielded a higher resolving reconstruction which was filtered using the Relion post-processing tool and sharpened with an automatically calculated B-factor of −73 Å^2^. The final resolution according to the FSC 0.143 criterion (Rosenthal and Henderson, 2003) is 2.96 Å (3.3 Å at FSC 0.5). The local resolution of both the poised state and active state reconstructions was assessed using ResMap (Kucukelbir et al., 2014) by supplying the unfiltered half-maps as inputs (Figure S3).

#### Model Building and Refinement

PDB crystal structures for the nucleosome (1kx3), isolated Dot1L residues 5-332 (1nw3), 4-330 (3qow) and ubiquitin (1ubq) were rigid-body fitted in the maps for the poised (3qow) and active state Dot1L (1nw3) using UCSF Chimera (Goddard et al., 2007). Next, both Dot1L-nucleosome structures were manually rebuilt using COOT (Emsley et al., 2010) and extended in several regions where the density allowed. Both models were subjected to alternating local and global real-space refinement using PHENIX (Adams et al., 2010) and COOT (Emsley et al., 2010). To assess for overfitting and avoid overinterpretation, we refined the models of poised and active state Dot1L against one of the sharpened, masked half-maps of the respective dataset (map_work_) and validated the structure against the second half-map (map_test_), similarly sharpened and masked. Specifically, the poised Dot1L reconstruction was refined using reference restraints from ubiquitin and Dot1L as well as secondary structure restraints to better account for the lower resolved Dot1L density. We also calculated individual atomic displacement parameters (ADP’s) by optimizing the real-space cross correlation between the model and the experimental map as implemented in phenix.real_space_refine (Afonine et al., 2018). The active Dot1l reconstruction was iteratively refined against the sharpened half-map applying alternating rounds of local and global refinements. Additional secondary structure restraints and reference restraints for the ubiquitin were applied and ADP’s were calculated. The final structures were validated against the second half-map of the respective reconstruction (map_test_) (Figure S3) and the FSC between the model and each half map (FSC_work_ and FSC_test_) and the full map (FSC_Full_) was determined. The computed FSC between the model and the second half-map (FSC_test_) agrees well with the FSC of the refined model against the target map used for refinement (FSC_work_). The resolution estimate of the model/map FSC_Full_ agrees well with the FSC 0.143 cutoff applied between the two half maps (Table 1 and Figure S3) and in all model/map FSC calculations, no significant correlation is measured that exceeds the calculated map resolutions. Both observations indicate that the built structures were not overfit nor overinterpreted versus their refinement targets. The final structures show excellent stereochemistry (see Table 1), as assessed by Molprobity (Chen et al., 2010).

